# A Comprehensive Methodological Framework for Anthropometric Head Shape Modeling Using Small Dataset

**DOI:** 10.1101/2024.06.02.597069

**Authors:** Leonardo H. Wei, Sudeesh Subramanian, Sajal Chakroborty, Suman Chowdhury

## Abstract

Detailed anthropometric characterization of complex shapes of human heads can ensure optimal fit, comfort, and effectiveness of head-mounted devices. However, there is a lack of a reliable and systematic approach for head shape classification and modeling for laboratory-based, small, occupation-specific datasets. Therefore, in this study, we proposed a streamlined framework comprising six steps—pre-processing, feature extraction, feature selection, clustering, shape modeling, and validation—for head shape classification and modeling. We collected 36 firefighter 3D head scans and implemented the framework. Different clustering techniques, such as k-means and k-medoids, were evaluated using the squared Euclidean distance of individual head shapes from their cluster centroid. Furthermore, five variations of NURBS and cubic spline methods were assessed to design the representative head shape of each cluster, and their accuracy was evaluated using mean square error (MSE) values. The clustering results indicated that k-means provide better metrics than k-medoids. Among the shape modeling methods, cubic spline least squares displayed the lowest MSE (0.70 cm^2^)and computational time (0.14 s), whereas NURBS least squares displayed the highest MSE (7.19 cm^2^). Overall, the framework with k-means clustering and cubic spline least squares modeling techniques proved to be the most efficient for small datasets.

## 1. Introduction

Anthropometric head shape modeling is essential for ensuring functional fit, safety, and the efficacy of head-mounted products (e.g., helmets, virtual/augmented reality, headgear, glasses, headphones, etc.)^1,2^, in addition to designing population-specific head-neck finite-element models^3^, head implant design^4,5^, and head and brain surgical planning^6,7^. Traditionally, one-dimensional (1D) head anthropometry measures (e.g., head breadth, length, circumference, etc.)—obtained manually using rulers, tape measures, and anthropometers^8^— have been utilized to develop head models because of their ease of data collection and processing. However, the manual measurement process can be time-consuming and error-prone if untrained personnel and uncalibrated equipment are used during the process. In contrast, with the advancement of scanning technologies, the use of three-dimensional (3D) body scanners has garnered much attention in the past two decades, as they can capture intricate geometrical variations of body shape modeling in a highly accurate manner^9^. Consequently, many recent 3D anthropometric and 3D head shape datasets have been developed in various countries by using 3D body scanning technologies^10–12^.

Although 3D body scanners are faster, less labor-intensive, and incomparably more accurate and detailed than traditional anthropometry, they produce a vast amount of 3D point clouds with complex geometries, curves, and contours. In comparison to 1D-based measures, these complex 3D scan datasets require additional data analysis steps, such as surface alignment, parameterization, and dimensionality reduction, before using for body shape modeling^1^. Such data processing stages are primarily attributed to between-subject variations in sitting behavior, cervical lordotic curvatures, and the challenges in maintaining the same reference coordinate system across the subjects during the scanning process. In order to mediate these challenges, previous studies suggested various techniques to align the 3D scan data in the same reference plane or coordinate system. Some of these techniques include iterative closest point method^13^ to align the head shapes with a reference, selecting reference facial landmarks^14^, and/or matching the model anatomical planes with a global reference plane manually^15–17^. Additionally, since a 3D scanned shape contains a massive amount of dense 3D data points, some previous studies18-21 have utilized principal component analysis (PCA), a dimensionality reduction technique, to reduce data dimensionality and select major features that are significantly different from each other while protecting the original shape. All these intermediate steps make the 3D shape-based anthropometric head modeling more complex and method-intensive than 1D-based measures.

In addition, the complexity of 3D head shape modeling becomes further compounded by the wide variations in head shape between subjects, which can be attributed to factors such as age, gender, and ethnicity ^14,22^. As a result, large datasets have been utilized to develop various population-specific anthropometric head shapes. In those studies, clustering techniques such as k-medoid^19–21^, hierarchical clustering^15,23,24^, and k-means clustering^25–27^ were commonly used to cluster the most similar head shapes in groups. However, there were putative results in the utility of k-means and k-medoids in 3D shape clustering. For instance, Lacko, et al.^19^ performed three variations of k-medoids—anthropometric-based, shape-based, and constrained-shape-based— in a set of 100 head scans. They found that three clusters provided the optimal number of clusters and that the shape-based clustering methods resulted in less variation within each cluster. Using this k-medoid method, Huang, et al. ^21^ clustered the 339 Chinese heads and reported that seven clusters were necessary. In another study, Niu, et al.^26^ employed k-means and clustered 378 Chinese military personnel heads, which also found that seven clusters were required. To extend the accuracy comparison of clustering procedures, Zhang, et al. ^28^ employed k-means, hierarchical, and fuzzy clustering techniques on the principal components of the points representing the head and reported that six clusters using k-means provided the highest accuracy. In summary, there is no consensus on using k-means or k-medoids and determining the optimal number of clusters, which can be subjective and application-specific dependent ^26^. Techniques such as Ray-Turi index^29^, silhouette coefficient^30^, and elbow method^31^ have been developed in an effort to determine the optimal number of clusters. However, these methods do not provide any statistical significance when selecting the optimal number of clusters.

When using clustering algorithms, a typical approach is to use the cluster centroid as the cluster representative shape. This approach may be effective for large datasets with sufficient scope for removing outliers. However, this approach is inappropriate for shape modeling with small datasets, as removing outliers can introduce skewness^32^. In an effort to overcome the limitations of clustering, recent studies have employed statistical shape modeling ^33–35^ to predict representative shapes based on statistical learning. This approach also requires large datasets to achieve reasonable accuracy. However, collecting 3D head scans of large populations poses significant challenges in laboratory-based academic setups owing to logistical hurdles and financial constraints. Thus, head shape modeling using a small dataset often becomes an inevitable option in academic laboratories, making it essential for researchers to identify appropriate clustering and head shape modeling techniques to ensure the accuracy of the available data and obtain meaningful results. In order to design customized head shapes for individuals, specialized curve fitting techniques, such as non-uniform rational B-splines (NURBS)^36^ and cubic splines ^37^, have been employed. These techniques allow for shape editing and customization, making them ideal for reconstructing complex shapes or designing products to meet individual anthropometric needs^38–42^. Customized shape products can indeed provide an enhanced fit, but they come with the drawback of being more expensive and time-consuming to fabricate. Manufacturing customized products can be burdensome and costly, even within a small dataset. Therefore, clustering techniques in small datasets can improve shape prediction by constraining population characteristics. In this context, NURBS and cubic splines techniques could potentially reduce the effects of outliers and improve the prediction accuracy of a cluster representative shape. So far, these techniques have not yet been attempted in conjunction with clustering procedures.

Therefore, this study aimed to determine accurate and efficient methods for clustering and modeling anthropometric head shapes while using a small dataset. While achieving this overarching goal, we come across a few methodological innovations: 1) develop a principal component analysis (PCA) shape-based clustering using k-means and k-medoids methods and determine the most accurate technique for clustering small datasets, 2) propose a method based on ANOVA for identifying the optimal number of clusters that yield statistical significance, and 3) employ NURBS and cubic spline curve fitting techniques to create a representative head shape of each cluster in the context of small datasets.

Furthermore, anthropometric shape models are required to develop head-mounted products that properly fit the head anatomical details, especially for first responders such as firefighters, law enforcement officers, and army personnel. Particularly, firefighters are heavier than the general population due to the demands of their job^43,44^, which require wearing heavier protective equipment and the ability to carry/drag people out of danger zones. A product developed using a civilian dataset may not be effective for occupation-specific populations, such as firefighters, law enforcement officers, and soldiers, who require unique anthropometric characteristics to meet their job-specific physical demands^45^. Yet, to date, the majority of these 3D anthropometric databases are based on civilian populations, like the Civilian American and European Surface Anthropometry Resource (CAESAR)^46^, National Institute of Occupational Safety and Health^47^, size China^48^, and Taiwanese population^49^. Furthermore, these databases were developed decades ago and may no longer be adequate for developing head-mounted devices for the new generations due to changes in their anthropometric characteristics ^50,51^. These shortfalls indicate a critical need for an updated firefighter 3D head shape dataset, particularly for designing their head protective devices.

## 2. Methodology

### 2.1 Experiment

#### Participants

At first, we scanned a sample of 36 firefighters’ heads (18 males and 18 females). However, the 3D scan data of seven females were discarded due to challenges in pre-processing their ponytails and uneven hair volume. Therefore, an additional seven males were recruited to acquire high-quality 3D scan data with minimal or no hair interference. This resulted in a participant cohort of 26 males (age = 37.6 ± 8.28 years; weight = 90 ± 17.17 kg; height = 177 ± 7 cm; BMI = 28.72 ± 4.58) and 10 females (age = 31.28 ± 7.17 years; weight = 63.11 ± 8.27 kg; height = 162 ± 5 cm; BMI = 24.28 ± 3.82). The participant inclusion criteria were that all participants were required to be free from any musculoskeletal, degenerative, or neurological disorders. Participants who met the inclusion criteria were asked to read and sign a consent form approved by the local institutional review board (IRB No: 2020-708).

#### Data acquisition

A laser-based handheld 3D scanner (EinScan HX, Shining 3D, Hangzhou, China) with two cameras, 0.04 mm accuracy, and 50 fps scanning speed was used for scanning participants’ head and neck complex (from T4 vertebral level to the tip of the head) in a controlled laboratory environment. All participants wore a swimming cap to minimize interferences from their hair volume. Female participants were specially instructed to fix their ponytail underneath the cap during scanning. In addition, all participants were asked to close their eyes and maintain their upright sitting posture during the 3D scanning process. Participants were instructed to remove any metallic or reflective materials, such as jewelry, to prevent interference with the laser scan. The scanning procedure was part of a broader project, which included three different scan conditions: no helmet, European jet-style helmet, and traditional helmet. Each scanning procedure took approximately 5 minutes, totaling 15 minutes for each participant’s scan. However, only the head scans without any helmet were used to assess anthropometry.

### 2.2 Methodological Framework for 3D head shape modeling

In order to sample and model the 3D head scan data from a relatively small participant pool, we provide a framework (Fig. 1) from a methodological perspective. The framework consisted of six major steps: 1) data pre-processing, 2) feature generation, 3) feature selection, 4) clustering, 5) modeling and prediction, and 6) validation. In the subsequent sections, we demonstrate the intricate details of these steps.

**Figure 1:**
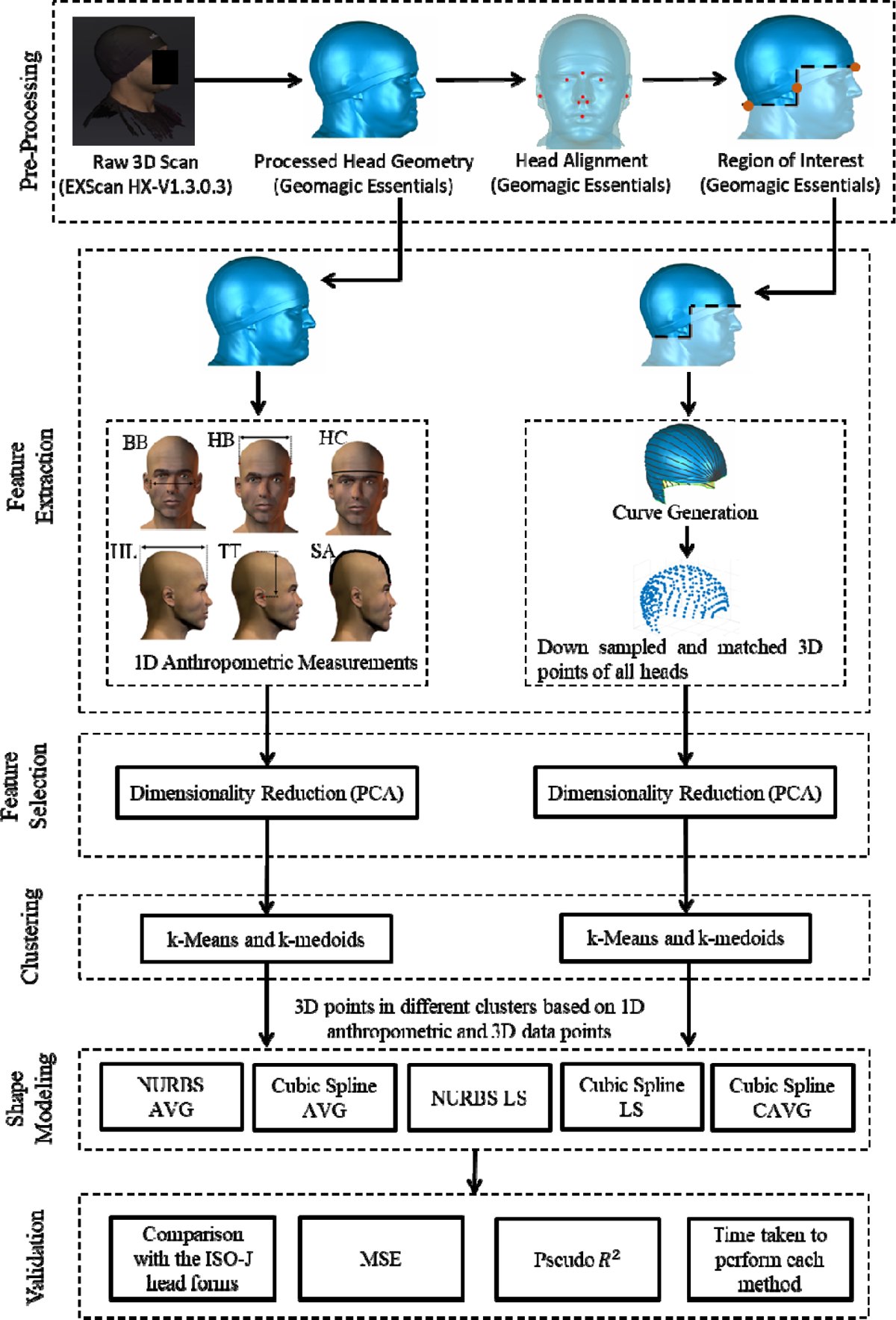
Methodological framework implemented in this study. In the shape modeling step, NURBS AVG and NURBS LS denote the application of non-uniform rational B-splines to the averaged data and non-uniform rational B-splines using least squares approximation, respectively. Cubic Spline AVG, Cubic Spline LS, and Cubic Spline CAVG respectively indicate the application of cubic splines to the averaged data, cubic splines using least squares approximation, and cubic splines obtained by averaging the coefficients of the cubic polynomials

#### 2.2.1 Data pre-processing

Our data pre-processing steps included 1) noise, artifacts, and unwanted body feature removal, 2) 3D surface alignment, 3) data smoothing, and 4) head region of interest (ROI) selection. As raw 3D scan data contains noises and unwanted surface details (outliers), we used EXScan HX-V1.3.0.3 (Shining 3D, Hangzhou, China) software platform to remove noises, unwanted body segments (shoulder, upper back, and sternum regions) and other artifacts, such as clothes from the 3D raw scan data (Figure 1). As no consistent reference axis was maintained during the 3D scanning process across the subjects, all scan data must be aligned on a consistent axis for downstream processing. We used the digital NIOSH large head form as a baseline head shape ^52^ and nine anthropometric landmarks (tip of the nose, right eye, left eye, mid-mouth, glabella, right ear opening, left ear opening, right nostril, and left nostril) to align 3D head shapes of all subjects to a reference axis (NIOSH). We used Geomagic Essentials software (Shining 3D, Hangzhou, China) to perform an automatic global registration process by considering those nine landmarks as constraints and thus aligned all scanned head shapes with the baseline reference head shape. We used an in-built smoothing function in Geomagic to smooth the 3D shape data and then cropped out unwanted neck, face, and skull segments to create the 3D head skull/scalp ROI for head shape clustering and modeling. The ROI was selected by intersecting three planes: a transverse plane passing through the glabella height, a sagittal plane passing through the tragion, and another transverse plane passing through the C1 height. The ROI comprised the upper region area resulting from the overlapping zones formed by the intersection of these planes (Fig.1).

#### 2.2.2 Feature Generation

As the literature shows, head shape modeling is based on both 1-D linear measurement and 3D scan point clouds, so we adopted both paradigms in this study to generate two sets of head features. The head features based on 1D anthropometric measurement were bizygomatic breadth, head breadth, head circumference, head length, tragion – top of the head, and sagittal arc. We measured these 1-D features in accordance with the definitions specified in the ANSUR II database^53^. As the 3D head shape (ROI) contained a vast amount of surface points (more than 10^6^ data points), which increases the complexity and efficiency of head shape clustering and modeling, we implemented a data reduction technique that created a total of 18 equally spaced head curves along the anterior-posterior (A-P) direction of individual head shapes. These curves were generated by creating a Frankfurt plane that rotated around an axis passing through the glabella and occiput landmarks at every 10° intervals, starting from the left tragion (0°) and ending at the right tragion (180°) points. We then generated 20 equally spaced surface points along the A-P direction at each one of the 18 curves of an individual head in Geomagic software. Thus, our second set of head features representing 3D head shape was created and contained a total of 360 equally spaced surface points for individual head surfaces.

#### 2.2.3 Feature Selection

The 1D anthropometric measures, hereinafter referred to as 1D measures, were initially normalized using the z-score to maintain a consistent scale across the measures. On the other hand, the 3D points were aligned to a common reference system, eliminating the need for normalization. Subsequently, principal component analysis (PCA) was conducted on both the normalized 1D anthropometric measures and the magnitude (Euclidean norm) of the 3D points to identify the most significant features and improve clustering accuracy^18^. The ideal number of principal components was determined by establishing a criterion of 90% of the total cumulative variance.

We performed Pearson Correlation analysis to relate the natural variation in head shape with the core anthropometric measures (body height, body weight, and BMI) and to compare the relationship between the selected head anthropometric measures. A Pearson Correlation value of more than 0.7, between 0.4 and 0.7, and less than 0.4 denoted strong, medium, and poor correlation between any two measures, respectively. These thresholds of *r* values were set based on a previous study developed by Akoglu ^54^.

#### 2.2.4 Clustering

We applied k-means^55^ and k-medoids^56^ on the full set of normalized 1D measures and on the extracted principal components of 1D anthropometric measures and 3D points to perform the clustering. To determine the optimal number of clusters, we repeatedly employed the clustering methods by varying the number of clusters. Each time the clustering was performed with a different number of clusters, the distance values of each cluster centroid from their respective subjects were calculated. It is important to note that the distance values were not directly calculated from the measures of 1D anthropometric measures or the spatial coordinates of 3D points. Instead, they were calculated based on the reduced dimensional space of principal components of 1D anthropometric measures and the magnitude of the 3D points. Subsequently, an analysis of variance (ANOVA) test was conducted to determine if the addition of an extra cluster based on the distance values displayed significant differences from the previous number of clusters (Figure 2). All significant tests were performed at a 95% confidence level (

**Figure 2:**
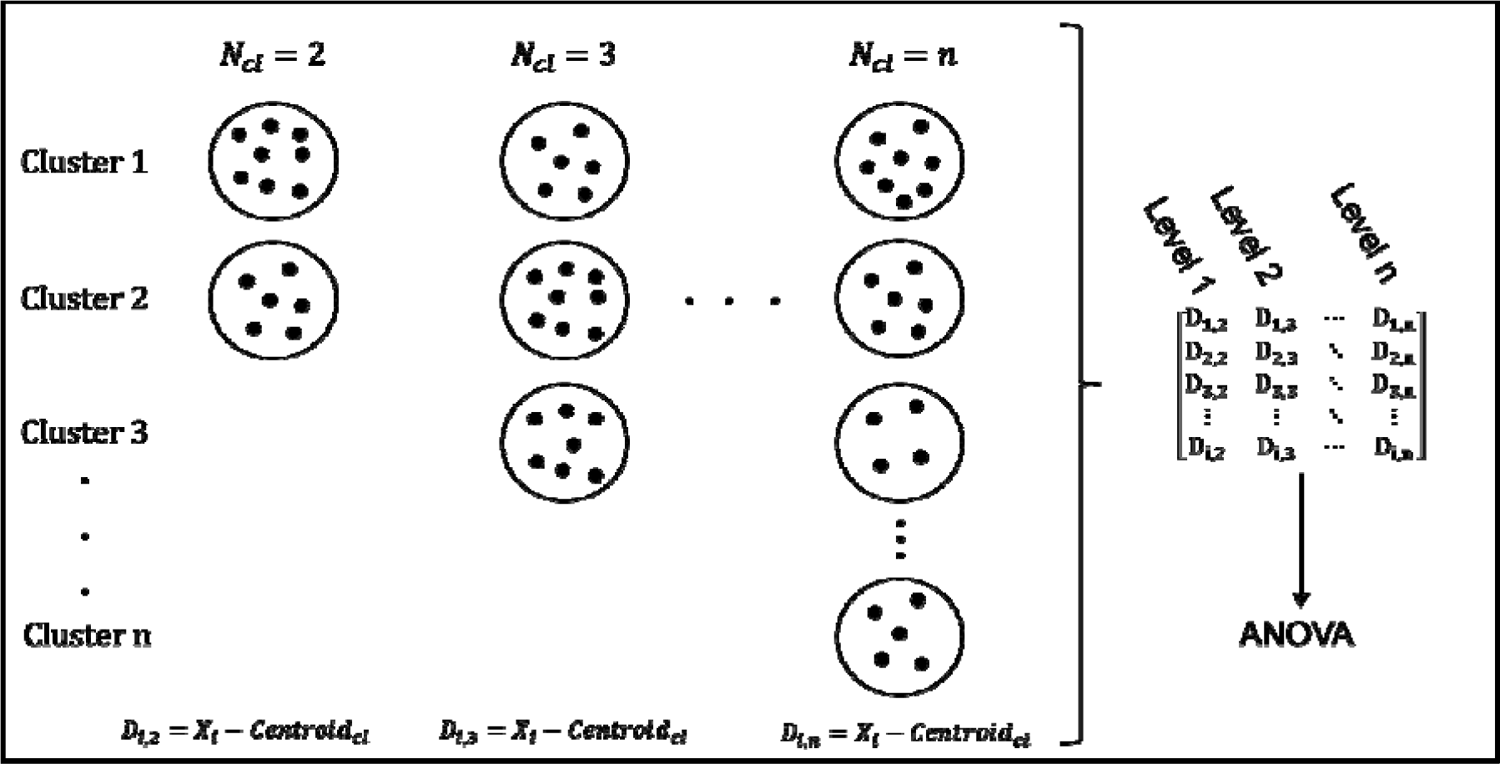
Workflow diagram to calculate the optimal number of clusters using ANOVA. The index “i” represents the subject number, the index “n” represents the total number of clusters and the index “cl”, the cluster number.

#### 2.2.5 Modeling and prediction

Curve fitting techniques are valuable tools to understand patterns, reduce noise, and highlight the most important characteristics in a given dataset. Methods such as linear regression^57^, polynomial regression^58^, cubic splines^36^, and B-splines^36^ are commonly used methods to fit data points. In particular, cubic splines and B-splines are the most used methods in modeling complex shapes due to their versatility in adjusting and modifying the curvature of shapes. Therefore, to model the head shapes of each cluster, non-uniform rational B-splines (NURBS) and cubic splines were chosen. For instance, the NURBS method provides local control over the shape of the curve through control points, allowing for local modifications without substantially affecting the global curve shape. On the other hand, cubic splines are defined by knot points that control the curvature of the shape. These knot points do not provide as much curvature flexibility as the control points in the NURBS technique, but they are simpler to implement and require less computational power.

Two main approaches— interpolation and approximation— were used to fit the NURBS and cubic splines to a given set of data points. The interpolation approach fits a function that passes exactly through all the given data points. In contrast, the approximation fits a function that does not pass through all the data points but provides a best-fit representation based on the available points. The current study will employ the interpolation approach on the resultant averaged head points for each cluster. In contrast, the approximation will consider each cluster’s entire dataset of head points. Additionally, a third approach was implemented by applying the cubic spline interpolation to each individual head point in a given cluster and then averaging these cubic spline coefficients across the subjects. The averaged coefficients were used to create the representative cubic spline function of the head in that given cluster. To facilitate notation, the interpolation methods were named non-uniform rational B-splines applied to the AVERAGE (NURBS AVG) and cubic spline applied to the average (Cubic Spline AVG), whereas the approximation methods were non-uniform rational B-splines least squares (NURBS LS) and cubic spline least squares (Cubic Spline LS). The coefficient averaging interpolation was labeled as Cubic Spline CAVG. Each one of these approaches will be further detailed in the following sections.

### NURBS AVG

A point in a NURBS curve (Eq. 2) is a weighted sum of control points, which is defined in the parameter space (*u*) and basis functions (N)^36^:

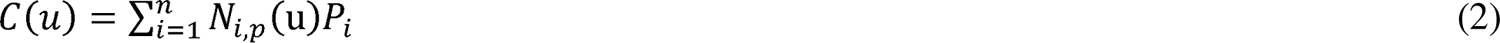

Here, *u* is a vector that contains the parameterized representation of the points, *P_i_* denotes the i^th^ control point, p is the degree of the spline, and *n* is the total number of control points. The control points are calculated based on pre-defined points manually selected in the interpolation process. The basis function was computed using the following recursive relation function^36^:

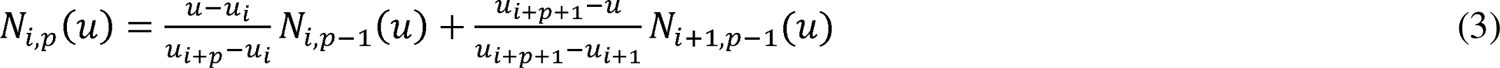

Note that:

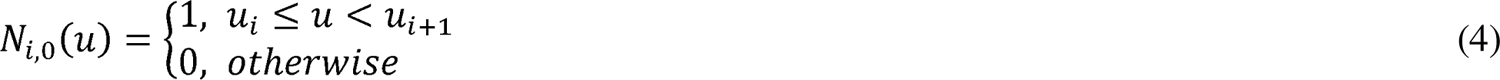

For more details, check algorithm 1 in the supplementary information.

### NURBS LS

In the least squares approach, the control point was selected by minimizing the following squared distance:

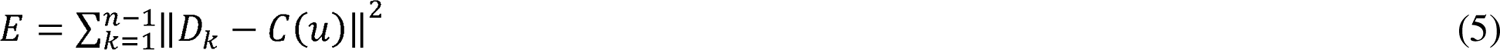

where *n* represents the total number of 3D points, *D*_k_ denotes the k^th^ 3D point, and *C(u)* is the NURBS curve representation. This curve representation, however, is slightly different from the previous standard NURBS curve:

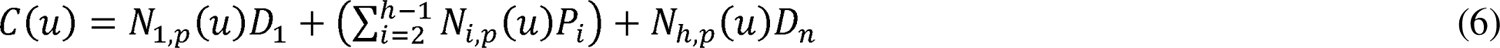

where, h, p, *D*_l_, *D_n_* and *P_i_* denote the number of control points, the degree of the spline, the first point, the last point, and the i^th^ control point, respectively. Six control points were selected for both methods, and their selection criteria were followed by a previous study ^59^. For more details, check algorithm 2 in the supplementary information.

#### Cubic Spline AVG

The cubic spline is a representation of continuous segments of cubic polynomials, which are separated by specific points called knots. In each of these knots, a new polynomial segment starts with different coefficients:

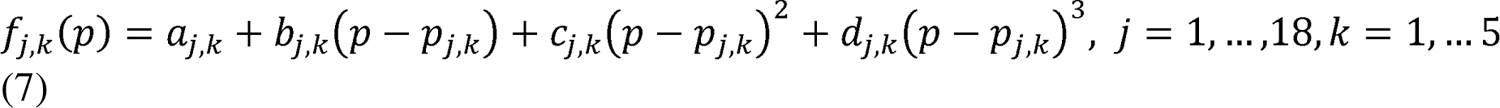

where *a_j,k_, b_j,k_, c_j,k_*, and *d_j,k_* are the coefficients of k^th^ segment in the j^th^ curve and *p_jk_* is the points to fit the cubic models. In the cubic spline AVG, these knots were averaged across subjects within a cluster, and the cubic spline model was applied to the averaged data points for each curve.

#### Cubic Spline LS

An independent parameterized variable (t) was created that represents the 3D points. This parameter space was normalized to range between 0 to 1 to account for the same scale across different subjects. The variable t is represented as:

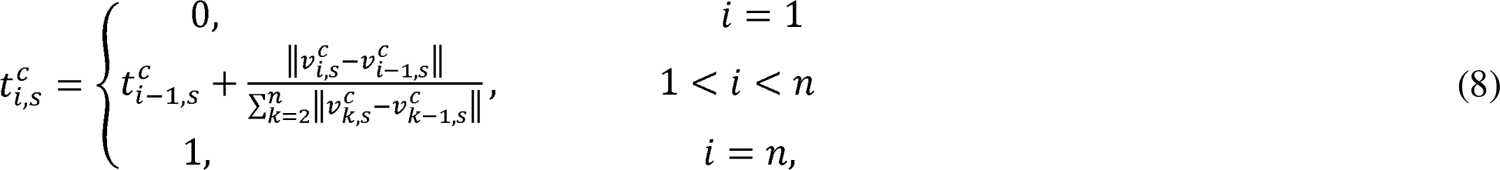

where *v_i_*_i_denotes the points (*x_i_, y_i_, z_i_*) for a given subject s for a given curve c. For each head curve, the variable *t^c^ _i,s_* will be a vector of size 20*ns* x 1, with *ns* representing the number of subjects in a given cluster. Thus, for each curve across all subjects in a given cluster, the minimization formula will be written as:

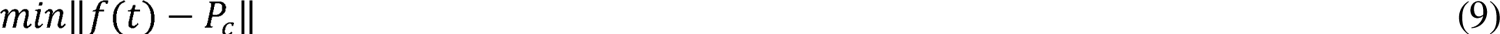

The variable *P_c_* represents the point coordinate in the direction c that ranges in x, y, and z, and *f(t)* represents the spline function applied in the parameterized variable t. For more details, check algorithm 3 in the supplementary information.

#### Cubic Spline CAVG

In the coefficient averaging method, the cubic spline model was fitted in the selected knots in each individual head for each cluster. After the model was fitted, corresponding coefficients were averaged across subjects (Eq. 10) within the same cluster:

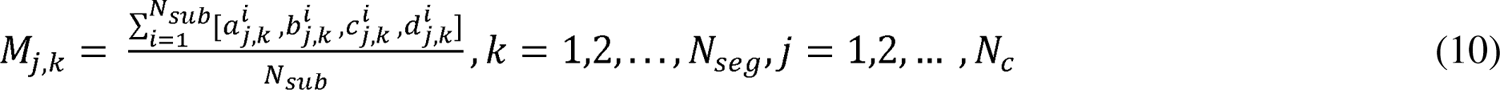

The subscripts *i*, *j*, and *k* are indices for subjects, number of curves, and number of segments, respectively. The term *M_j,k_* is a *N_c_* x 4*N_seg_* matrix of averaged coefficients, wherein each row comprises the averaged coefficients throughout one curve and each column, one coefficient.

#### 2.2.6 Validation

For the comparative analysis of head measures, four databases— ANSUR II ^60^, CAESAR ^61^, NIOSH^47^, and US Army^47,62^— were chosen. Descriptive statistics, such as the mean, standard deviation, minimum and maximum values, and the 5th and 95th percentiles of each head measure, were calculated. These statistics were then compared with each individual database as well as with the pooled average across all databases. To comply with standard nomenclature and for comparison, we initially compared and analyzed our cluster’s similarity with the five NIOSH headform ^63^. Subsequently, we classified our head forms based on standard terminology, such as small, medium, and large sizes.

To assess the accuracy of the model in predicting the head shape of each cluster, mean squared error (MSE) was performed to compare the prediction accuracy of each cluster:

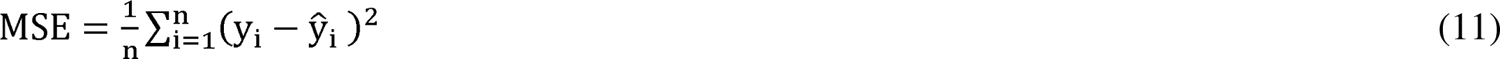

where *n* represents the number of data points, *y_i_* is the subjects’ 3D points, and *y_i_* is the predicted values from the models. A total mean squared error was also calculated to assess the overall accuracy of the methods across clusters.

The variability in the data was explained by calculating a pseudo r-squared (R^2^) for each individual method (Eq. 12). The pseudo r-squared was employed due to the non-linearity of the curves used in the fitting procedure.

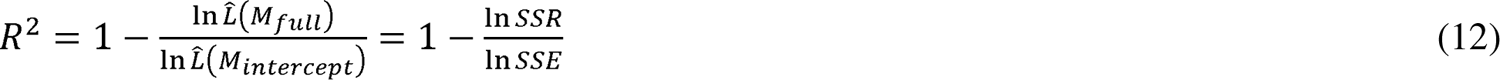

where, SSR is the sum squared of residuals, and SSE is the sum of squared error. Unlike the traditional r-squared, the pseudo r-squared typically yields lower values, ranging from 0.2 to 0.4 for a good fit ^64^. In addition, as the methods may achieve similar MSE and *R*^2^ scores, the run time to perform each method was also calculated to contribute to the determination of the most effective method.

The most effective curve fitting and clustering techniques were selected to generate the representative head shapes. These head shapes and curves were compared against the 3D head scans from the NIOSH ^47^ database using a 3D deviation analysis tool in Geomagic Essentials. Despite being a civilian-spanned database, it was the only available set that provided open-access 3D scan head shapes.

## 3. Results

### 3.1 1D-based head anthropometric measures

Our participants’ mean head anthropometric measures, BB, HB, HC, and HL, were higher by 3.3%, 7.2%, 4.1%, and 1%, respectively, compared to the pooled averages of the reference datasets (Table 1). Additionally, the minimum BB (13.15 cm), HC (54.73 cm), HL (18.84 cm), and TT (11.2 cm) of our dataset were 14.35%, 15.22%, 23.95%, and 6.7% higher than the minimum value found in other databases. Only the minimum head breadth was found to be lower than other databases due to an outlier.

**Table 1.** Comparison of descriptive statistics of anthropometric measures of the participants with reference datasets. Except for Caesar, other reference datasets did not measure all anthropometric measures, and, therefore, those missing measures are omitted from this comparison. All dimensions are in cm.

|  | This Study | ANSUR II <sup>60</sup> | CAESAR <sup>61</sup> | NIOSH<sup>47</sup> | US Army<sup>47,62</sup> | Pooled data |
| --- | --- | --- | --- | --- | --- | --- |
| **Bizygomatic Breadth** |  |  |  |  |  |  |
| 5<sup>th</sup> Percentile | 13.17 | 12.80 | 12.97 | 12.93 | 12.78 | 12.84 |
| 95<sup>th</sup> Percentile | 17.06 | 15.20 | 15.87 | 15.16 | 14.54 | 15.22 |
| Mean ± STD | 14.48 ± 0.99 | 13.97 ± 0.73 | 14.42 ± 0.88 | 14.04 ± 0.68 | 13.66 ± 0.54 | 14.02 ± 0.72 |
| Min ~ Max | 13.15 ~ 17.40 | 11.60 ~ 17.40 | 11.90 ~ 17.7 | 11.5 ~ 17 | 11.7 ~ 16.1 | 11.50 ~ 17.7 |
| **Head Breadth** |  |  |  |  |  |  |
| 5<sup>th</sup> Percentile | 14.56 | 14.20 | 14.00 | 14.11 | 14.05 | 14.09 |
| 95<sup>th</sup> Percentile | 18.10 | 16.20 | 16.28 | 16.04 | 15.78 | 16.11 |
| Mean ± STD | 16.19 ± 1.07 | 15.22 ± 0.62 | 15.14 ± 0.70 | 15.07 ± 0.59 | 14.92 ± 0.52 | 15.10 ± 0.61 |
| Min ~ Max | 11.68 ~ 18.18 | 13.10 ~ 18.00 | 13.30 ~ 17.20 | 12.9 ~ 17.9 | 12.6 ~ 17.3 | 12.6 ~ 18.00 |
| **Head Circumference** |  |  |  |  |  |  |
| 5<sup>th</sup> Percentile | 55.75 | 54.00 | 53.36 | 53.96 | 53.29 | 53.66 |
| 95<sup>th</sup> Percentile | 63.07 | 60.00 | 60.19 | 59.67 | 58.18 | 59.61 |
| Mean ± STD | 58.95 ± 1.72 | 57.00 ± 1.83 | 56.78 ± 2.08 | 56.81 ± 1.74 | 55.74 ± 1.48 | 56.63 ± 1.81 |
| Min ~ Max | 54.73 ~ 63.55 | 50.00 ~ 63.50 | 51.10 ~ 62.20 | 47.5 ~ 65.4 | 50 ~ 62.7 | 47.50 ~ 65.40 |
| **Head Length** |  |  |  |  |  |  |
| 5<sup>th</sup> Percentile | 19.04 | 18.20 | 20.83 | 18.17 | 18.08 | 18.75 |
| 95<sup>th</sup> Percentile | 21.36 | 21.00 | 25.05 | 20.58 | 20.16 | 21.75 |
| Mean ± STD | 20.44 ± 0.68 | 19.63 ± 0.85 | 22.94 ± 1.28 | 19.37 ± 0.73 | 19.12 ± 0.63 | 20.25 ± 0.91 |
| Min ~ Max | 18.84 ~ 21.67 | 16.80 ~ 22.50 | 19.39 ~ 27.46 | 15.20 ~ 22.5 | 15.8 ~ 22 | 15.20 ~ 27.46 |
| **Tragion Top** |  |  |  |  |  |  |
| 5<sup>th</sup> Percentile | 11.85 | 11.80 | 12.92 | -- | -- | 12.3 |
| 95<sup>th</sup> Percentile | 14.82 | 14.00 | 15.28 | -- | -- | 14.58 |
| Mean ± STD | 13.18 ± 0.80 | 12.96 ± 0.67 | 14.10 ± 0.72 | -- | -- | 13.44 ± 0.69 |
| Min ~ Max | 11.2 ~ 14.92 | 10.50 ~ 15.00 | 11.87 ~ 16.22 | -- | -- | 10.50 ~ 16.22 |
| **Sagittal arc** |  |  |  |  |  |  |
| 5<sup>th</sup> Percentile | 38.22 | -- | 27.95 | -- | -- | 27.95 |
| 95<sup>th</sup> Percentile | 42.34 | -- | 33.32 | -- | -- | 33.32 |
| Mean ± STD | 40.25 ± 1.30 | -- | 30.63 ± 1.63 | -- | -- | 30.63 ± 1.63 |
| Min ~ Max | 35.98 ~ 43.28 | -- | 25.82 ~ 35.45 | -- | -- | 25.82 ~ 35.45 |

When comparing individual databases, only the mean TT and HL of CAESAR were found to be higher than our dataset. The descriptive statistics of the sagittal arc in our study were substantially larger than those reported in CAESAR, the only other study that provided this measure. This discrepancy is attributed to differences in the sagittal arc measurement methodology. In our study, it was measured from the glabella point to the C0-C1 joint (complete skull arc), whereas in CAESAR, it was measured from the glabella to the occiput bone.

Overall, a higher mean and range of our anthropometric measures indicate that our small firefighter dataset comprises mainly medium to large head sizes. After comprehensively comparing all measures with the individual reference datasets, we found that our dataset was most closely aligned with ANSUR II, followed by CAESAR and NIOSH data. As ANSUR II and CAESAR did not provide a 3D digital head form, we only used the NIOSH dataset for further analysis and comparison.

The Pearson correlation analysis between head anthropometric measures and body mass, height, and BMI revealed that tragion-top and sagittal arc were the least correlated variables (Figure 3). Additionally, it was observed that head anthropometric measures are not strongly correlated with subject height, mass, and BMI. On the other hand, strong correlations (coefficients greater than 0.7^54^) were observed between head circumference and bizygomatic breadth and between head length and head circumference.

**Figure 3:**
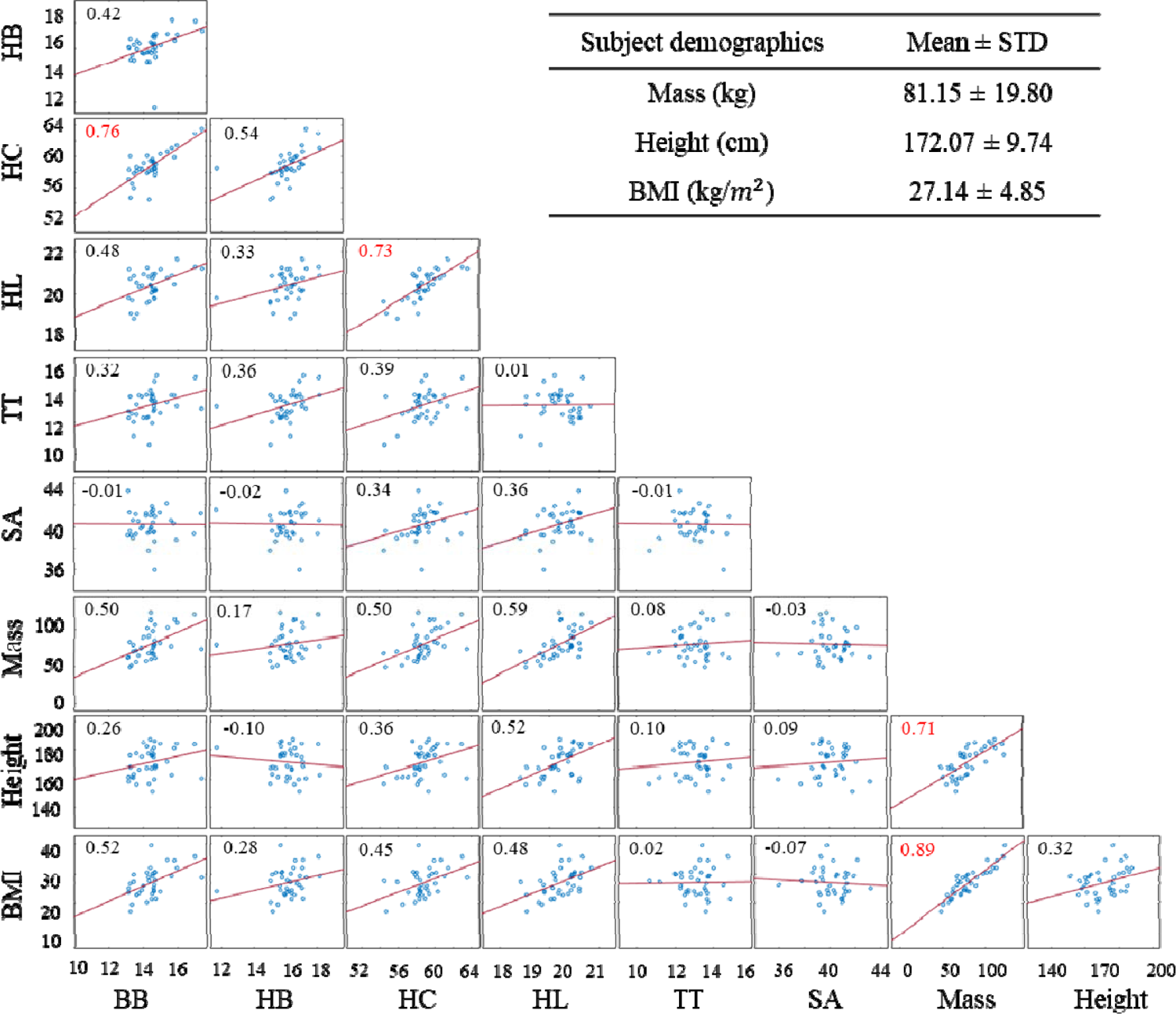
The Pearson correlation matrix illustrating the relationships among the head anthropometric measures and mass, height, and BMI. In each box’s left top corner, correlation values are depicted, representing the correlation coefficient between pairs of variables. Strong correlations are highlighted in red. Abbreviations: BB, bizygomatic breadth; HB, head breadth; HC, head circumference; HL, head length; TT, tragion top; SA, sagittal arc; BMI, body mass index.

**Figure 4:**
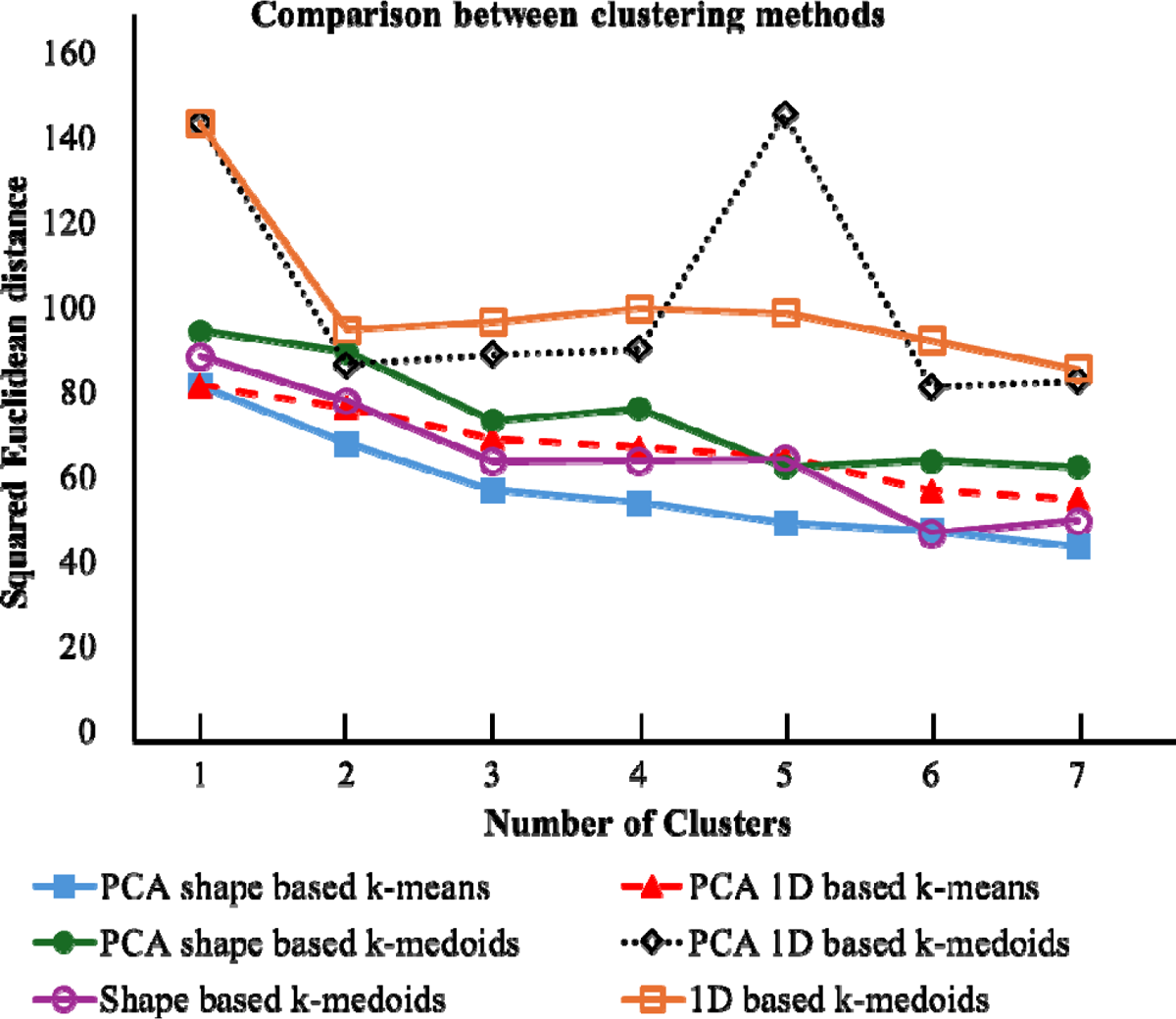
Squared Euclidean distance comparison between different k-means and k-medoids clustering approaches.

The principal component analysis revealed that four principal components for 1D anthropometric measures and seven principal components for 3D shape-based covered more than 90% of the variance in the data. The first principal component (PC 1) of 1D anthropometric measures accounted for more than 56% of the total variance (Supplementary data 1; Figure 1) in the data. Therefore, those 1D measures that provided the most weightage in this PC (head circumference, bizygomatic breadth, and head length) were considered for head shape sizing performed after the clustering process (Supplementary data 1; Table 1).

### 3.2 Clustering

The PCA shape-based k-means displayed the lowest squared Euclidean distances among all the clustering methods, closely followed by the shape-based k-medoids and PCA 1D-based k-means. On the other hand, PCA 1D-based k-medoids and 1D-based k-medoids displayed the highest squared Euclidean distances among all methods, with the PCA 1D-based k-medoids showing slightly lower values than the 1D-based k-medoids. Interestingly, the PCA shape-based k-medoids displayed the highest values than the shape-based k-medoids. To compare the differences between shape-based methods and traditional 1D anthropometric measures, the 1D-based k-means and PCA shape-based k-means were chosen due to the lowest squared Euclidean distances within their respective categories (1D- and shape-based). Our ANOVA tests indicated that three clusters were enough to display significant differences (p=0.03) between each other for 1D anthropometry-based clustering. In contrast, four clusters were required for the PCA shape-based k-means clustering (p=0.04).

Clustered head shapes are presented in Fig. 5. In 1D anthropometry-based clustering, Cluster 1 (two males and five females) comprised small head sizes (BB: 13.74 cm (average), HC: 57.01 cm, and HL: 19.49 cm) among all clusters, while cluster 2 (three males and one female) contained the largest (BB: 16.51 cm, HC: 62.26 cm, and HL: 21.28 cm) head size (Fig. 5a). Cluster 3 (20 males and five females) covered medium head size (BB: 14.37 cm (average), HC: 58.96 cm, and HL: 20.57 cm). The same trend was observed for Cluster 1, 2, and 3 of 3D shape-based clustering, but with slight differences in cluster distribution. In the 3D shape-based clustering, Cluster 1 (BB: 13.41 cm (average), HC: 57.09 cm, and HL: 20.07 cm) consisted of one male and six females, Cluster 2 (BB: 15.93 cm, HC: 61.47 cm, and HL: 21.11 cm) included three males, and Cluster 3 (BB: 14.49, HC: 58.89 cm, and HL: 20.47 cm) comprised 13 males and three females. An additional cluster, Cluster 4 (BB: 14.79 cm, HC: 59.58 cm, and HL: 20.44 cm), is exclusively present in the 3D clustering method and is composed of seven males and three females. This 4^th^ cluster seemed to have a slightly higher HC than Cluster 3, with a comparatively larger occipital region (Fig. 5b).

**Figure 5:**
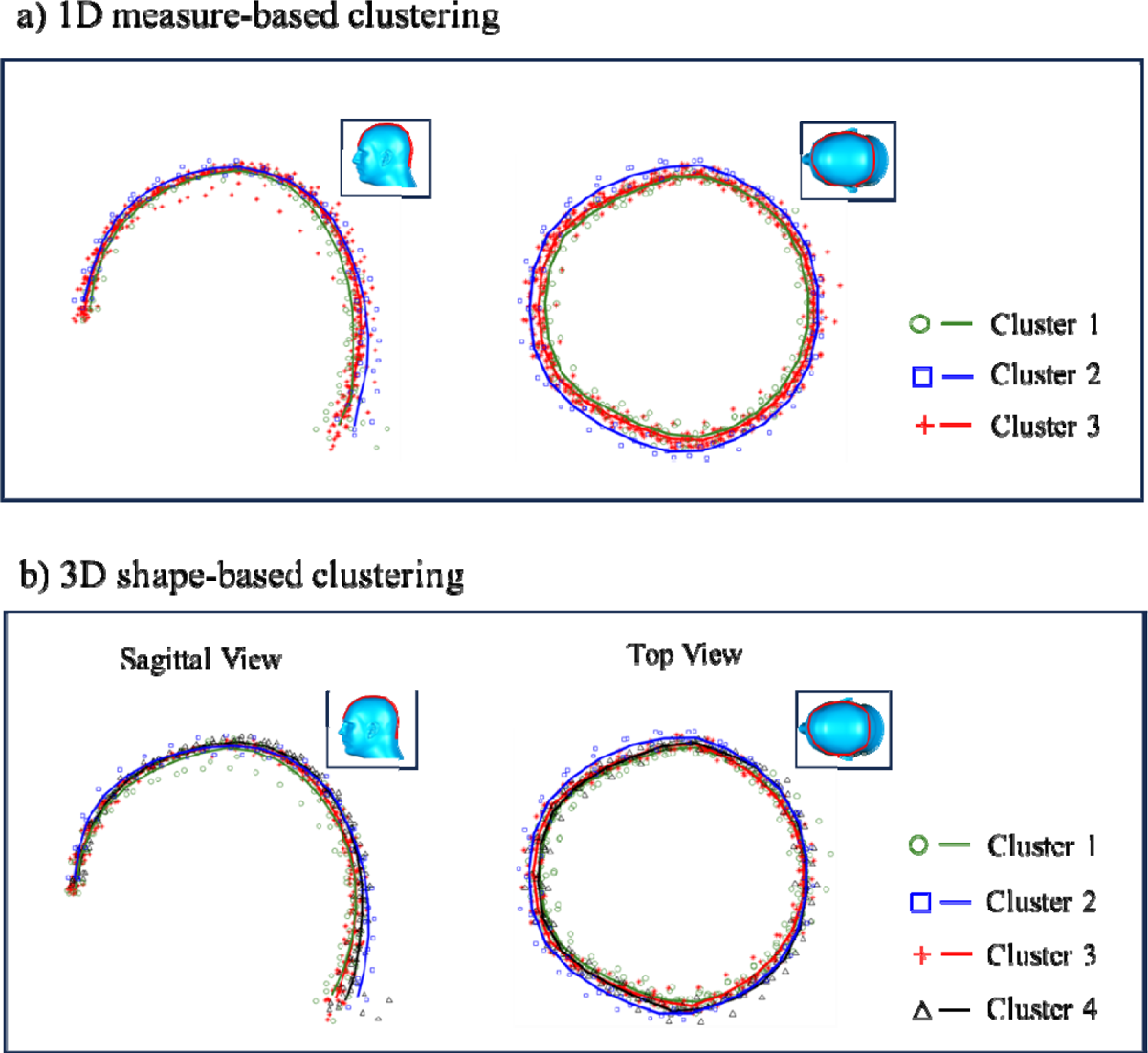
Sagittal and top views of different clusters for a) 1D anthropometric-based clustering and b) 3D shape-based clustering. Markers indicate subjects’ 3D head points, while solid lines indicate the cluster average.

Compared with the NIOSH head forms, our clusters were medium to extra-large head sizes (Table 2). In 1D anthropometric-based clustering, Cluster 1’s head anthropometric measures were slightly higher than those in NIOSH medium-size head form. Cluster 2 was larger than the NIOSH large head form. Except for BB and TT, all other head anthropometric measures in Cluster 3 were found to be higher than the NIOSH large head form. Similarly, in 3D shape-based clustering, Cluster 1 was slightly larger than the NIOSH medium head form, although TT was slightly lower. Cluster 2 had all head anthropometric measures larger than the NIOSH large head form, while Cluster 3 and 4 had higher HB, HC, HL, and SA. Overall, Clusters 2, 3, and 4 were closer to the NIOSH large head form. In terms of overall size, Cluster 2 was the largest, followed by Cluster 4, Cluster 3, and Cluster 1.

**Table 2:** A comparison of average head anthropometric measures (in centimeters) of each cluster from 1D anthropometric measure based and 3D point based methods and with NIOSH data. Abbreviations: BB, bizygomatic breadth; HB, head breadth; HC, head circumference; HL, head length; TT, tragion top; SA, sagittal arc.

|  |  | Mean<br>(min ~ max) |  |  |  |  |  | Closest<br>Matching<br>NIOSH<br>Headform | Size<br>Classification |
| --- | --- | --- | --- | --- | --- | --- | --- | --- | --- |
|  |  | BB | HB | HC | HL | TT | SA |  |  |
| 1D | Cluster 1<br>(n 25) | 13.74<br>(13.6 ~<br>14.7) | 16.00<br>(15.22<br>~<br>16.87) | 57.01<br>(54.73<br>~<br>58.57) | 19.49<br>(18.84<br>~<br>20.14) | 13.02<br>(11.20<br>~<br>14.58) | 38.68<br>(35.98 ~<br>39.59) | Medium | Medium |
|  | Cluster 2<br>(n=7) | 16.51<br>(15.66<br>~<br>17.40) | 17.66<br>(17.06<br>~<br>18.18) | 62.26<br>(61.08<br>~<br>63.55) | 21.28<br>(20.90<br>~<br>21.27) | 13.61<br>(12.87<br>~<br>14.80) | 39.96<br>(38.75 ~<br>41.24) | Large | Extra Large |
|  | Cluster 3<br>(n=4) | 14.37<br>(13.17<br>~<br>15.81) | 16.01<br>(11.68<br>~<br>17.11) | 58.96<br>(57.63<br>~<br>60.65) | 20.57<br>(19.56<br>~<br>21.27) | 13.15<br>(11.97<br>~<br>14.92) | 40.74<br>(39.39 ~<br>43.28) | Large | Large |
| 3D | Cluster 1<br>(n=16) | 13.41<br>(13.15<br>~<br>14.02) | 15.72<br>(15.17<br>~<br>16.30) | 57.09<br>(54.73<br>~<br>59.14) | 20.07<br>(18.84<br>~<br>21.20) | 12.39<br>(11.20<br>~<br>13.64) | 40.10<br>(38.92 ~<br>43.28) | Medium | Medium |
|  | Cluster 2<br>(n=10) | 15.92<br>(14.42<br>~<br>17.40) | 16.96<br>(16.50<br>~<br>17.31) | 61.47<br>(59.43<br>~<br>63.55) | 21.11<br>(20.43<br>~<br>21.67) | 13.31<br>(12.87<br>~<br>14.02) | 39.81<br>(38.75 ~<br>41.24) | Large | Double Extra<br>Large |
|  | Cluster 3<br>(n=7) | 14.49<br>(13.62<br>~<br>15.66) | 15.94<br>(11.68<br>~<br>18.18) | 58.89<br>(57.63<br>~<br>61.08) | 20.47<br>(19.56<br>~<br>21.27) | 13.29<br>(12.31<br>~<br>14.92) | 40.50<br>(38.62 ~<br>41.90) | Large | Large |
|  | Cluster 4<br>(n=3) | 14.79<br>(13.25<br>~<br>17.00) | 16.68<br>(15.69<br>~<br>18.09) | 59.59<br>(58.17<br>~<br>62.98) | 20.44<br>(19.08<br>~<br>21.31) | 13.50<br>(11.97<br>~<br>14.80) | 40.091<br>(35.98 ~<br>42.17) | Large | Extra Large |

Cluster 1 in both anthropometric measure-based and 3D shape-based clustering is larger than the NIOSH medium head form but smaller than the large head form, so we classified that as medium in this study. Cluster 3 of anthropometric measure-based and 3D shape-based clustering was classified as large. Cluster 4 of 3D shape-based clustering was classified as extra-large. Cluster 2 of the anthropometric measure-based clustering and 3D shape-based clustering was double-extra-large.

### 3.3 Shape modeling

The average MSE of cluster head shapes generated from 1D anthropometric-based clustering was higher than those generated using 3D head shape-based clustering (Table 3). The average MSE values for the latter case were 8.41% to 25% lower than the former case, except when the NURBS LS method was used. NURBS LS method showed 13.23% higher MSE values when applied to 3D shape-based clustering compared to the 1D anthropometric-based clustering. Between different shape modeling methods, Cubic Spline LS and Cubic Spline CAVG exhibited the highest accuracy (lowest MSE), closely followed by NURBS AVG. Cubic Spline AVG and NURBS LS were the least accurate and were significantly different from other methods.

**Table 3:** Mean square error of predicted shapes from 1D anthropometric-and 3D-based clustering (in cm^2^)

| 1D anthropometric-based clustering |  |  |  |  |  |
| --- | --- | --- | --- | --- | --- |
|  | Cluster 1<br>(n=25) | Cluster 2<br>(n=7) | Cluster 3<br>(n=4) | Average MSE |  |
| NURBS AVG | 0.81 ± 0.87 | 1.67 ± 1.46 | 0.74 ± 0.79 | 0.98 |  |
| NURBS LS | 5.31 ± 2.31 | 11.60 ± 6.03 | 3.66 ± 1.97 | 6.35 |  |
| Cubic Spline AVG | 2.13 ± 1.50 | 2.90 ± 1.85 | 2.12 ± 1.57 | 2.26 |  |
| Cubic Spline LS | 0.79 ± 0.84 | 1.62 ± 1.47 | 0.66 ± 0.58 | 0.94 |  |
| Cubic Spline CAVG | 0.79 ± 0.83 | 1.62 ± 1.51 | 0.65 ± 0.43 | 0.94 |  |
| Shape-based clustering |  |  |  |  |  |
|  | Cluster 1<br>(n=16) | Cluster 2<br>(n=10) | Cluster 3<br>(n=7) | Cluster 4<br>(n=3) | Average<br>MSE |
| NURBS AVG | 0.62 ± 0.66 | 0.51 ± 0.35 | 0.54 ± 0.46 | 1.34 ± 1.12 | 0.75 |
| NURBS LS | 8.60 ± 2.25 | 7.69 ± 2.38 | 7.18 ± 2.27 | 5.28 ± 2.28 | 7.19 |
| Cubic Spline AVG | 1.92 ± 1.30 | 1.80 ± 0.59 | 1.91 ± 1.05 | 2.67 ± 1.99 | 2.07 |
| Cubic Spline LS | 0.60 ± 0.59 | 0.50 ± 0.39 | 0.51 ± 0.40 | 1.19 ± 0.68 | 0.70 |
| Cubic Spline CAVG | 0.60 ± 0.59 | 0.50 ± 0.39 | 0.51 ± 0.40 | 1.19 ± 0.68 | 0.70 |

Pseudo-R-squared values (*R*^2^) of Cubic Spline CAVG and Cubic Spline LS had the highest values, explaining the variability in the data better than other methods, closely followed by NURBS AVG (Table 4). Despite the good *R*^2^, the computational time was substantially high for the Cubic Spline CAVG method (12.15s). On the other hand, Cubic Spline LS appeared to be the between different clustering techniques, 3D shape-based clustering yielded slightly better R^2^ scores. Therefore, Cubic Spline LS applied to 3D shape-based clustering was considered a better approach in this study.

**Table 4:** Pseudo *R*^2^and time taken to perform the regression for the 1D anthropometric measurement and 3D shape-based clustering methods.

| 1D Anthropometric measurements |  |  |  |  |  |  |
| --- | --- | --- | --- | --- | --- | --- |
|  | Cluster 1<br>(n=25) | Cluster 2<br>(n=7) | Cluster 3<br>(n=4) | Average | Time Taken (s) |  |
| NURBS AVG | 0.256 | 0.247 | 0.264 | 0.256 ± 0.008 | 0.95 |  |
| NURBS LS | 0.141 | 0.179 | 0.124 | 0.148 ± 0.028 | 1.37 |  |
| Cubic Spline AVG | 0.201 | 0.202 | 0.230 | 0.211 ± 0.017 | 0.19 |  |
| Cubic Spline LS | 0.260 | 0.249 | 0.263 | 0.257 ± 0.008 | 0.09 |  |
| Cubic Spline CAVG | 0.260 | 0.249 | 0.261 | 0.257 ± 0.006 | 12.15 |  |
| Shape-based clustering |  |  |  |  |  |  |
|  | Cluster 1<br>(n=15) | Cluster 2<br>(n=10) | Cluster 3<br>(n=7) | Cluster 4<br>(n=4) | Average | Time Taken (s) |
| NURBS AVG | 0.256 | 0.253 | 0.316 | 0.240 | 0.266 ± 0.034 | 1.54 |
| NURBS LS | 0.173 | 0.119 | 0.172 | 0.179 | 0.161 ± 0.028 | 1.87 |
| Cubic Spline AVG | 0.203 | 0.248 | 0.233 | 0.191 | 0.219 ± 0.026 | 0.26 |
| Cubic Spline LS | 0.265 | 0.246 | 0.320 | 0.244 | 0.269 ± 0.035 | 0.14 |
| Cubic Spline CAVG | 0.267 | 0.245 | 0.321 | 0.244 | 0.269 ± 0.036 | 18.28 |

### 3.4 Point-to-Point Deviation Analysis with NIOSH head form

A point-to-point deviation analysis with NIOSH head forms was then performed to compare with the representative head forms of 3D shape-based clusters generated using Cubic Spline LS (Fig. 6). Warmer colors (> 0 deviation) indicated that those regions in our head forms were smaller than the corresponding regions of the NIOSH head forms, while colder (< 0 deviation) colors indicate that those regions were larger in our head forms. The predominantly negative deviations of all our head forms from NIOSH’s small and short-wide head forms indicated that our dataset comprises larger sizes. Cluster 1 (medium) was found to have the least deviations with the NIOSH long narrow and medium head forms, whereas Cluster 2 (double extra-large) had substantial negative deviation (average: −0.41) even against the larger NIOSH head form, supporting our previous anthropometric-based comparison. The average positive deviation of Cluster 3 (large) indicated that it is slightly smaller than the large NIOSH head form. In Cluster 4 (extra-large), it can be observed that its occipital and frontal region is similar to NIOSH’s large head form, but the mid-section (left to right temporal region) was larger in size.

**Figure 6:**
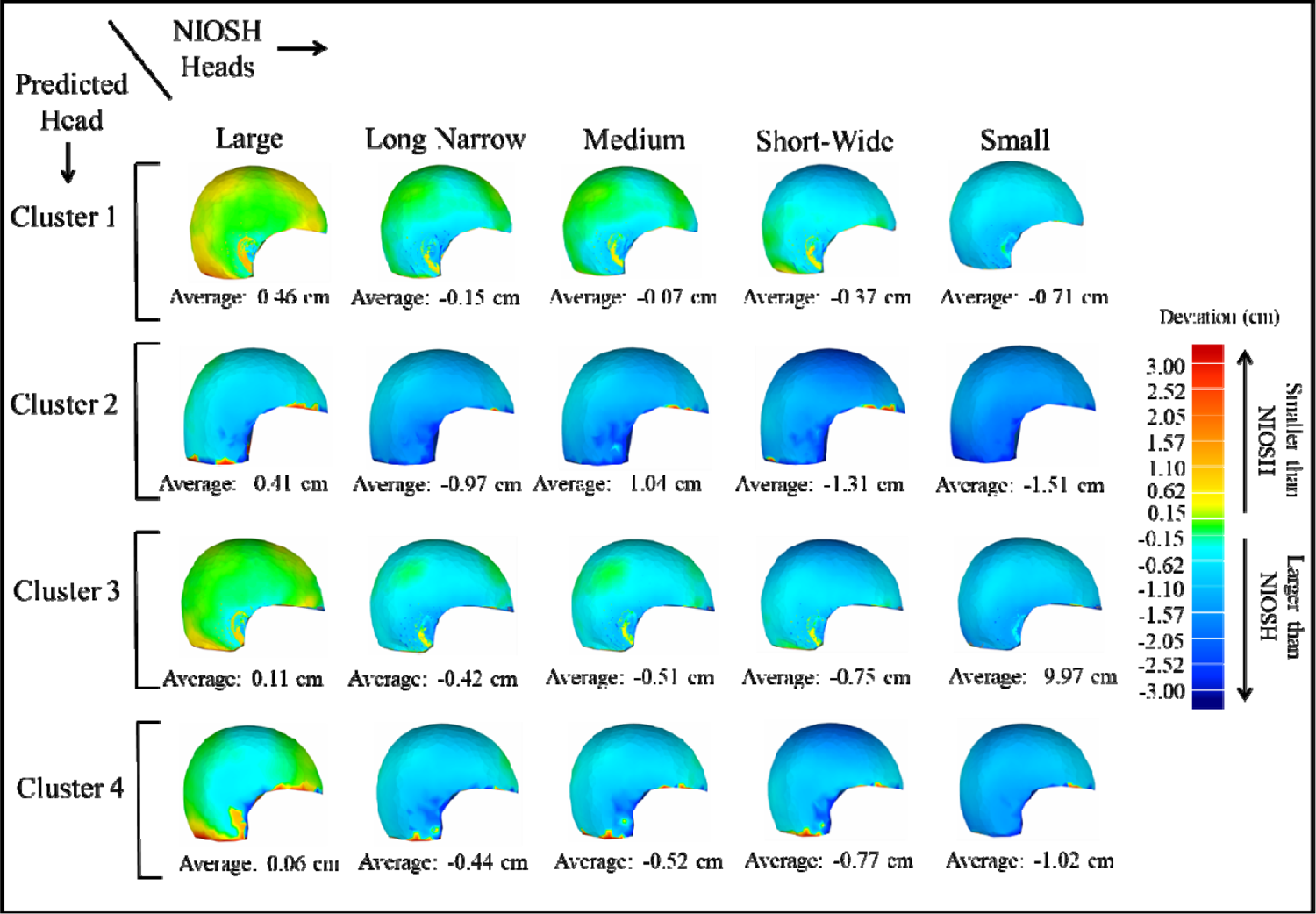
Comparison with the NIOSH 3D head forms by performing deviation analysis, with the average point-to-point distance and the standard deviation between the predicted head from the cubic spline least squares method and each one of the five NIOSH head forms.

## 4. Discussion

Head forms for developing head-mounted devices for firefighters and law enforcement officers are typically based on general population datasets. However, these specific populations have unique anthropometric characteristics that distinguish them from the general population. Still, even between these occupation-specific datasets, anthropometric characteristics can differ substantially due to many underlying factors related to job demands. For instance, in this study, the average head circumference, head breadth, and head length were found to be larger than in other studies^17,25,61,63^. Our data were closer to the most recent military personnel dataset (ANSUR II^60^), while most deviations were observed with the previous US military personnel dataset^62^ collected three decades ago. This discrepancy could be related to generational anthropometric changes due to improvements in nutrition and lifestyle^65^, indicating the need to update the anthropometric dataset, specifically for highly physically demanding occupational jobs like firefigters^66,67^. In addition, it was observed that overall, our dataset was larger than the CAESAR dataset, which comprises American and European civilian populations. This agrees with previous research^43,44,68^ that anthropometric characteristics of physically demanding occupations such as firefighters and law enforcement officers differ from general population characteristics. Firefighters are generally heavier when compared to the general population, with their weight at least 8 ∼ 10 kg higher than US civilians^43^. Even though our study did not show a strong correlation between BMI and head anthropometric measures, a previous study^69^ indicated that higher BMI was directly associated with larger head dimensions, such as head circumference. Therefore, these assessments indicate the necessity of collecting occupation-specific population datasets for the development of head-mounted devices.

This study’s results indicated that, in general, PCA enhanced the clustering efficiency. It could be argued that PCA would not be required for a small number of features. Nevertheless, it was observed that the squared Euclidean distances using k-medoids with the full set of measures (1D-based k-medoids) were higher than those using k-medoids on the reduced set of principal components (PCA 1D-based k-medoids), suggesting that PCA enhances clustering metrics even for a smaller number of features. While comparing the efficiency of k-means and k-medoids, the higher squared Euclidean distances observed in the k-medoids can be attributed to its method of selecting cluster centroids. Whereas in k-medoids, the centroid is an actual data point, in k-means, the centroid is calculated by averaging all points in a cluster. For small datasets, where even minor changes in data points can substantially affect the mean, the choice of using an actual data point as the centroid method could not adequately capture the variations. As the sample size increases, the distribution of the sample mean tends to be normally distributed ^70^, and the centroid of the k-medoids tends to be closer to the mean. For this reason, when dealing with large datasets, k-medoids are more suitable because computing the cluster centroid doesn’t require averaging all data points, thereby reducing computational time^71^. However, for small datasets, k-means is expected to perform better than the k-medoids^72,73^, which could be confirmed by the squared Euclidean distances reported in this study. As the k-means proved to be better for small datasets, 1D-based and shape-based inputs were selected to perform k-means and compare clustering results. The lower number of significant clusters reported in 1D-based clustering is due to the relatively less information provided by the 1D anthropometric measures. In contrast, shape-based clustering offers much more shape information by considering geometrical differences^20^, resulting in a larger number of clusters and better clustering accuracy^19^.

The highest MSE and low *R*^2^ values displayed by NURBS LS are related to the second-order continuity of NURBS curves, which guarantees higher curvature gradients but also implies higher oscillations^74^, consequently increasing the error. Opposing this trend, NURBS AVG MSE values displayed comparatively good MSE and *R*^2^values. The higher curvature and oscillatory nature of NURBS, in the averaging case, is beneficial, as averaging reduces the variability, which is counterbalanced by the higher oscillatory behavior of NURBS. Interestingly, applying the cubic spline to the average (Cubic Spline AVG) did not perform as good as NURBS applied to the average (NURBS AVG). This can be attributed to the minimal curvature of cubic splines when interpolating points ^75–77^. Despite all the limitations of cubic splines, Cubic Spline LS achieved better results than NURBS. For instance, In the least-squares approximation problem, all points are utilized instead of a reduced set of averaged points, and the fitted curve will incorporate more variability. However, the minimal curvature of cubic splines ^75^ can reduce this variability effect, improving the accuracy of this method. Achieving a similar accuracy as the Cubic Spline LS, the Cubic Spline CAVG is a good option to reduce the effects of outliers. By averaging the coefficients, the curvature of the resultant curve is softened, as points are not directly involved in the calculation. Nonetheless, this method requires more computational power. The NURBS method generally allows for more complex shape modeling when data is sparse due to its non-uniform knot selection ^78^. Furthermore, for design purposes, NURBS offers great flexibility for easily modifying control points’ position, which effectively changes the curvature of the shapes and enables shape edition. Nevertheless, the NURBS method requires a lot of parameters, such as knots and control points, which increases the computational power^79,80^. In that sense, cubic splines are easier and faster to implement and still provide good accuracy. If design edition is not required and computational power is limited, cubic spline least squares pose a good option due to its straightforward implementation and simplicity. However, if design edition and shape modification are required, NURBS would enhance the user experience and facilitate the creation of user-defined shapes.

Previous studies primarily used the cluster 3D centroid head points to provide representative heads for each cluster^17,21,24^ or used methods such as NURBS and cubic spline to fit geometries^37,40,81^ individually. However, predicting head shapes from smaller sample sizes can pose challenges since outliers may have a more pronounced impact in such scenarios. Relying solely on the cluster centroid could compromise the precision of head predictions in those cases. In contrast, our systematic framework employed clustering techniques alongside NURBS and cubic spline methods, demonstrating that this combination is effective for handling small sample sizes and provides an accurate solution for population-specific anthropometry.

There are some limitations that need to be acknowledged. First, the six anthropometric measures used in this paper were manually collected using Geomagic software. This approach is prone to human errors that could potentially assign subjects that do not actually belong to that cluster.

Secondly, it’s worth noting that as we employed the ear as the reference point for trimming the heads, some head scans exhibited longer occipital regions than others. This variation could influence the head shape classification in the 3-D shape-based clustering method, as it may assign heads to larger or smaller clusters, depending on the ear size. Thirdly, the heads were specifically trimmed in the region covered by the helmet, including the ear portion and the occipital region. Some details of the frontal region, such as the nose and eyes, could affect the clustering.

## 5. Conclusion

This study showed that conventional civilian anthropometry data is insufficient for designing head-mounted devices for occupation-specific populations. For smaller datasets, PCA-based k-means clustering outperformed k-medoids in accuracy for both 1D- and shape-based clustering, highlighting k-means’ superior performance with small sample sizes. Additionally, the higher accuracy and reduced computational time of the Cubic Spline Least Squares (LS) method suggested that capturing the variations within each cluster is more effective when utilizing the entire dataset rather than relying on the mean. The proposed methodological framework can potentially improve clustering and shape modeling accuracy, particularly for population-specific anthropometry, where the collection of large datasets may not be feasible.

## Acknowledgments

We would like to thank Felipe Zambrini Santos, Gustavo M. Paulon, Sneha Shrestha, and Kathryn Bell for their contributions to data processing and analysis.

## Authors Contribution

L.W. contributed to data collection and processing, analyzed the results, created figures, and primarily wrote the manuscript. S.S. co-analyzed and interpreted the results, generated figures, co-wrote and critically reviewed the manuscript. S.C. contributed to the presentation of mathematical formulations and critically reviewed the methodology and literature review. S.K.C. directed the study and analysis, obtained funding, and co-wrote and critically reviewed the article.

## Supplementary information

### Principle component analysis

**Supplementary Table S1:**
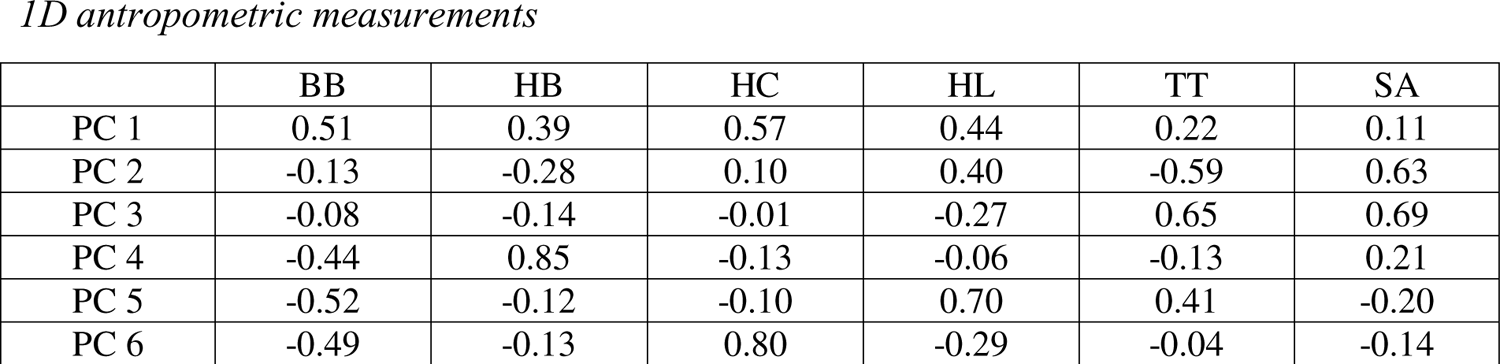
Coefficients of *1D* anthropometric measures for each principal component. Positive values indicate a positive correlation with the principal component and negative values indicate a negative correlation with the respective principal component.

### Algorithms

**Algorithm 1:**
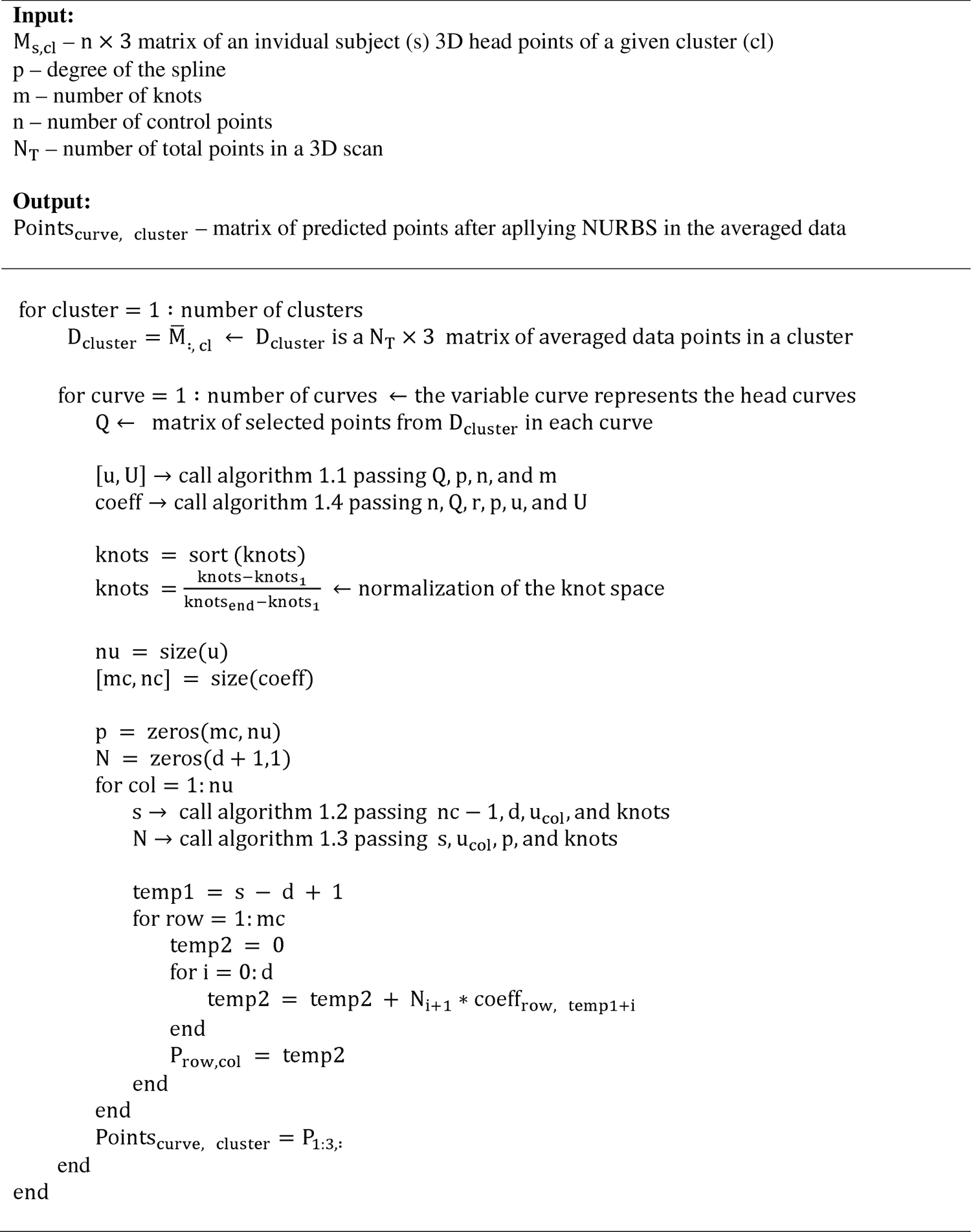
NURBS applied to the average (adapted from ^82^)

**Algorithm 1.1:**
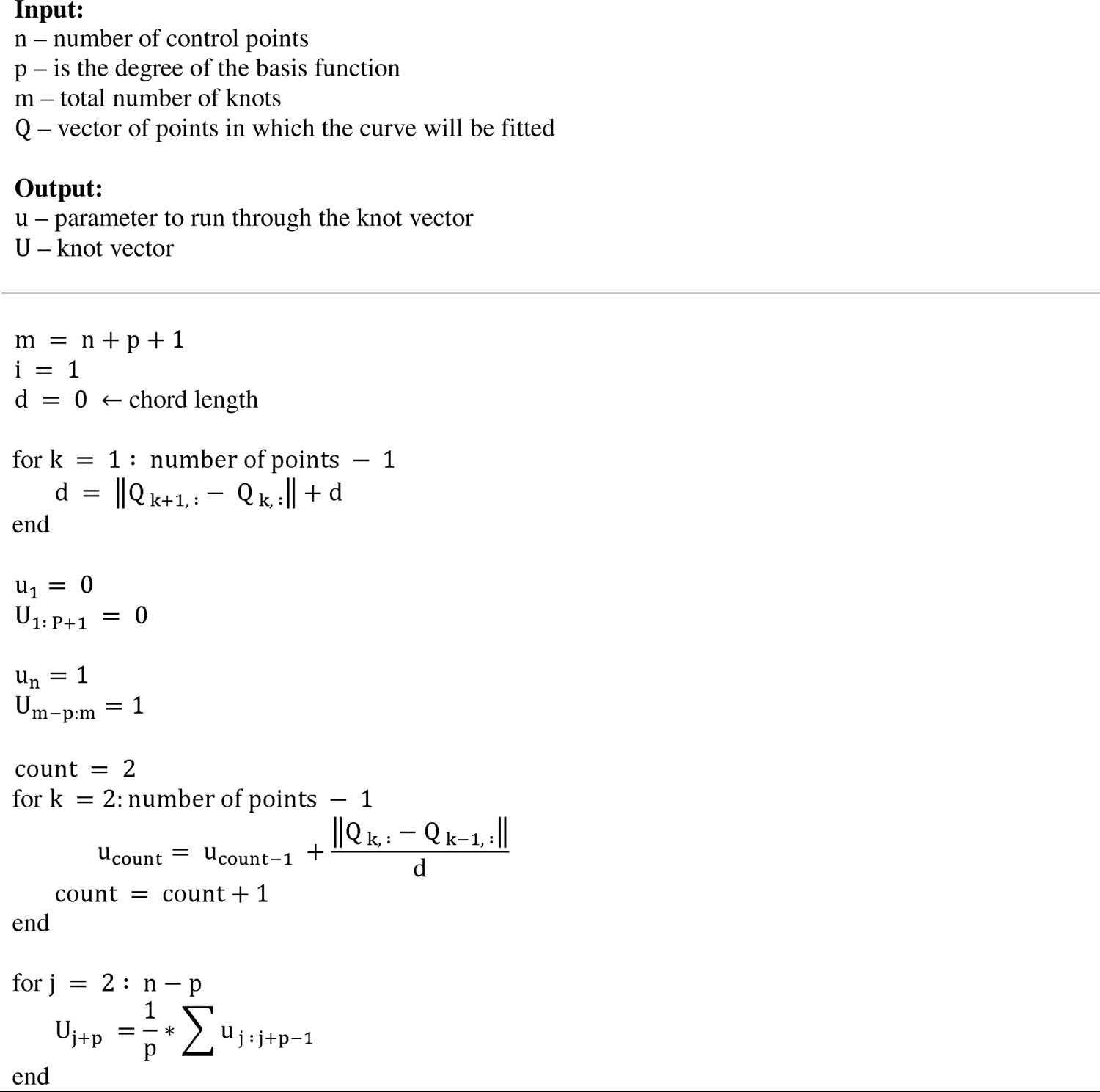
knot calculation.

**Algorithm 1.2:**
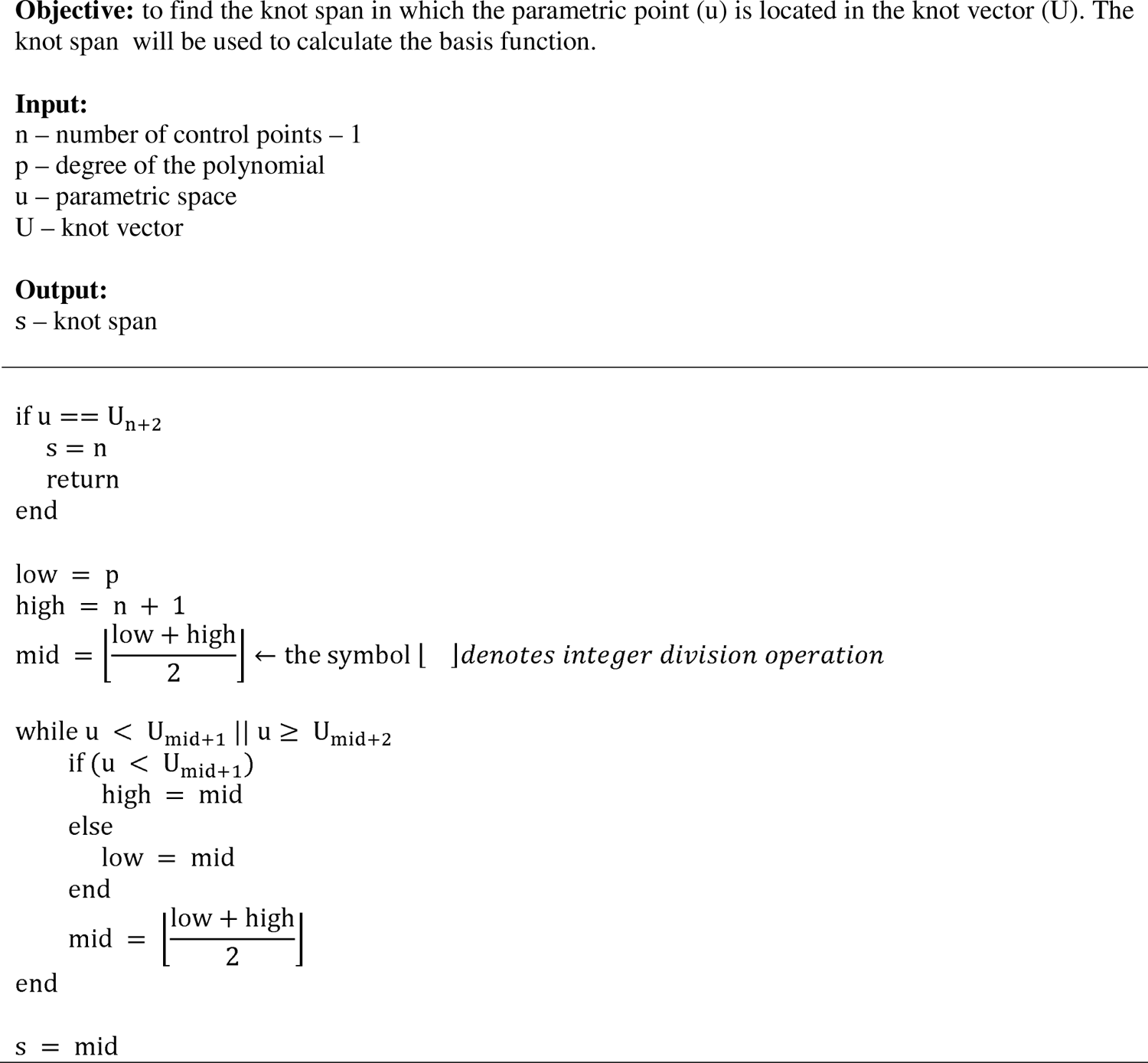
Find span

**Algorithm 1.3:**
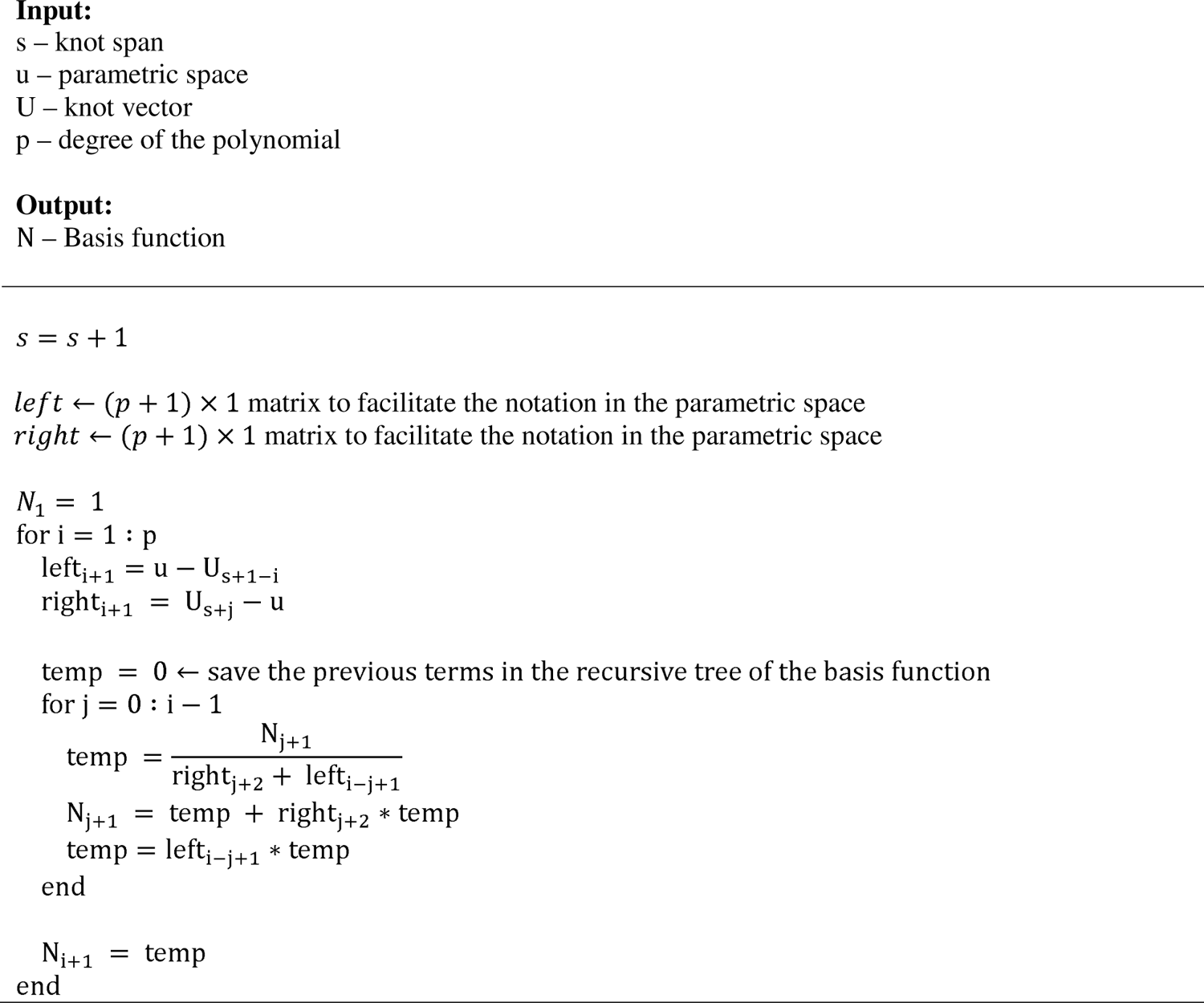
Basis function

**Algorithm 1.4:**
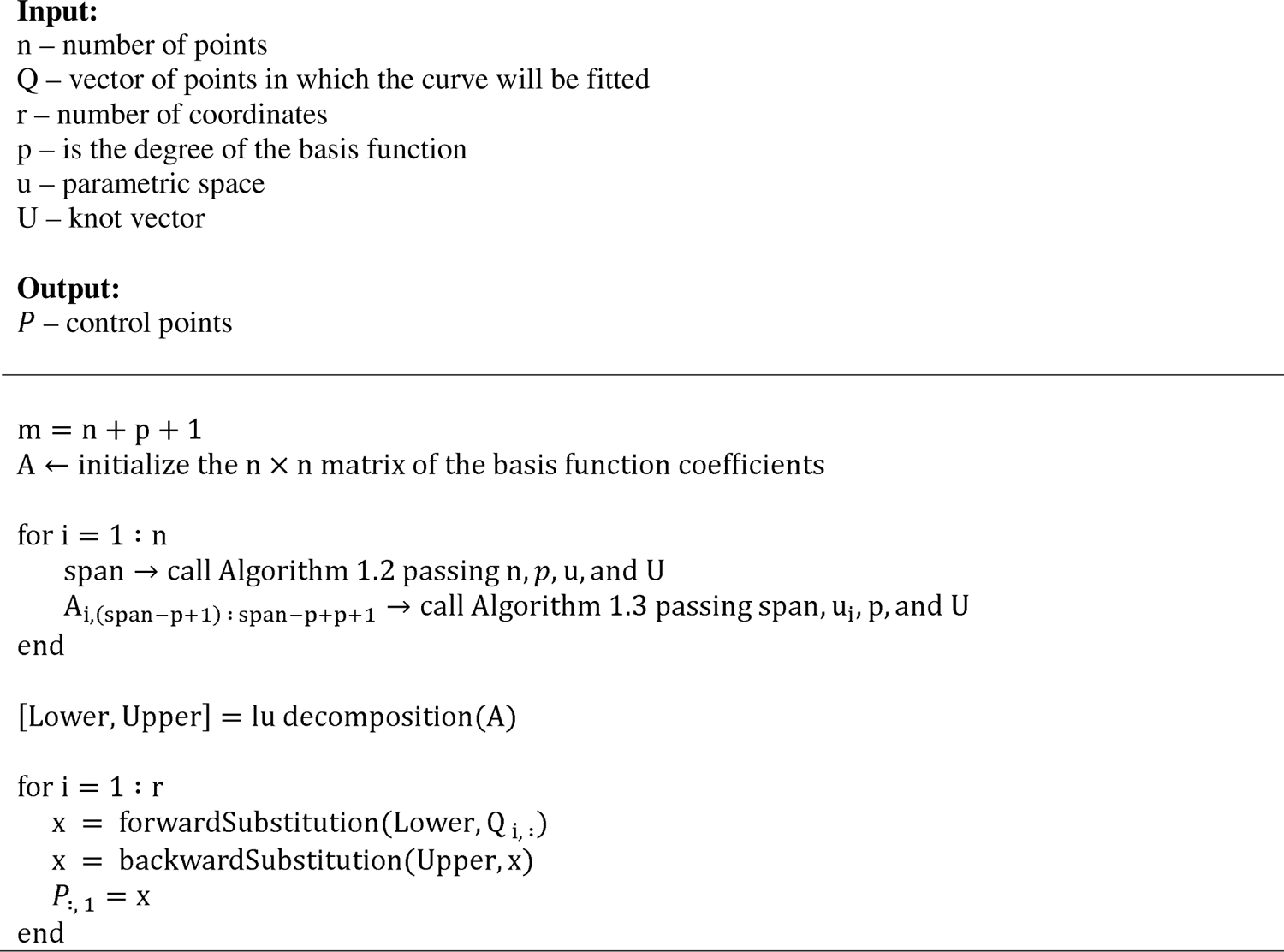
control points creation.

**Algorithm 2:**
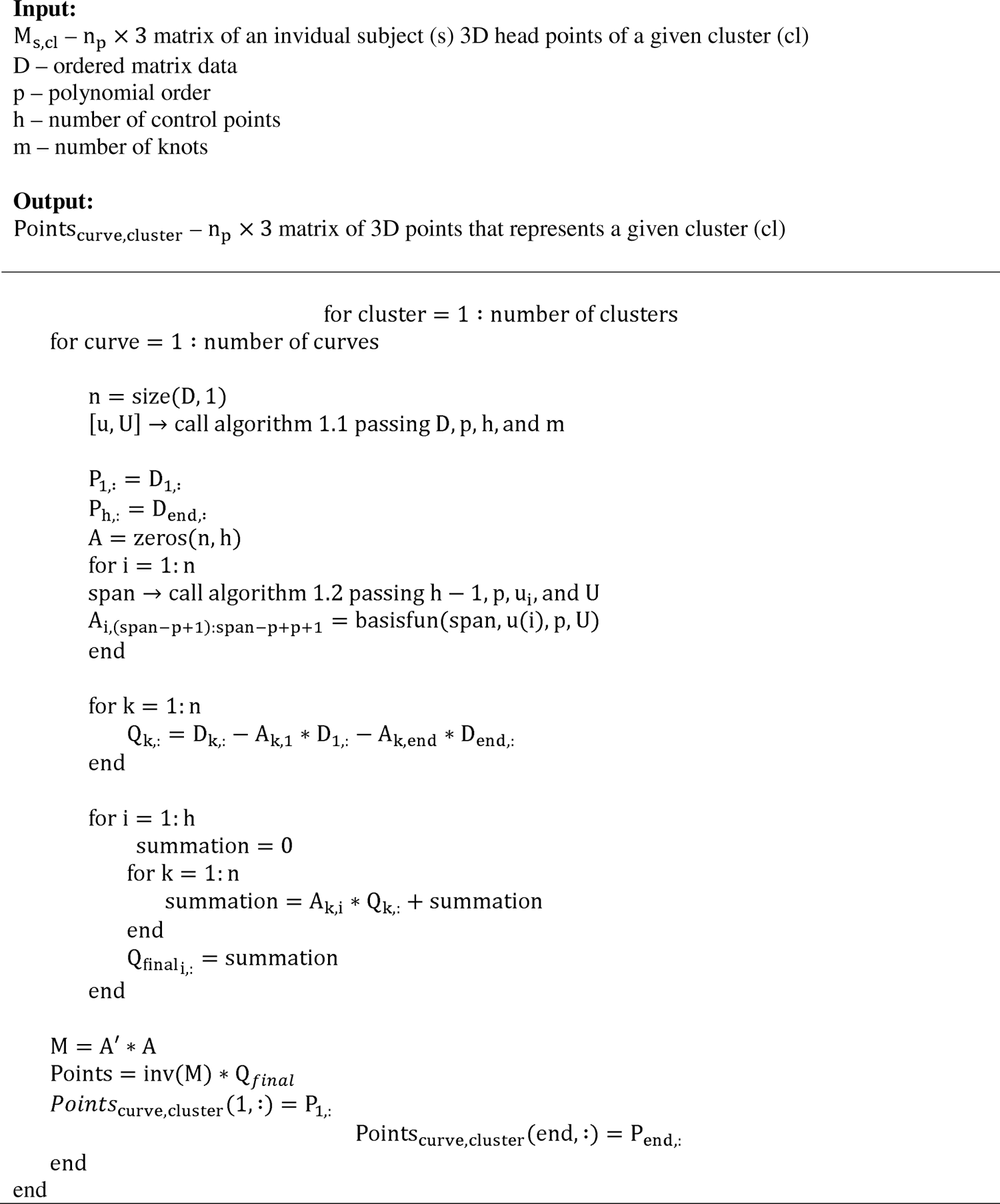
NURBS LS

**Algorithm 3:**
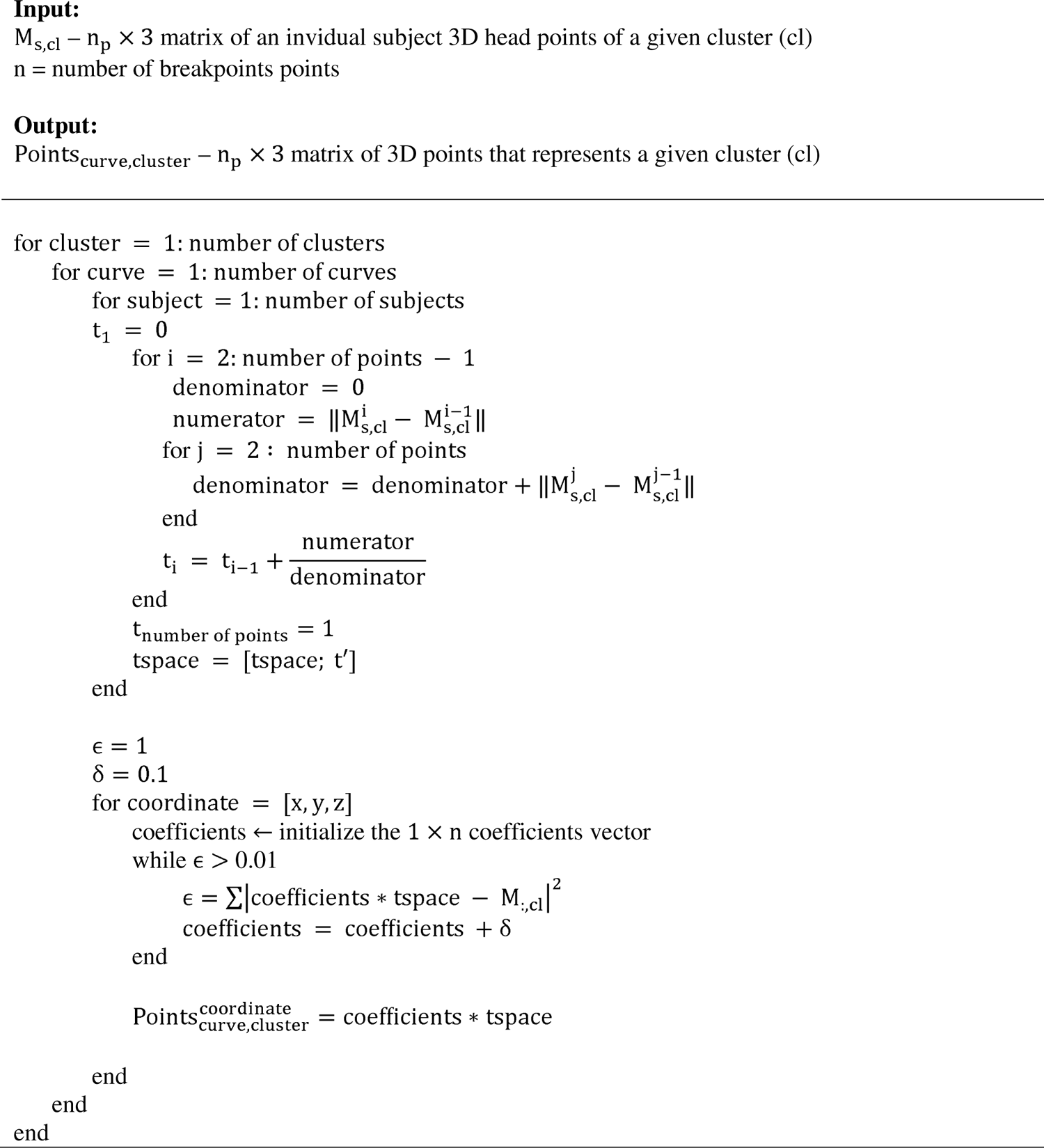
Cubic Spline LS

## References

1 Ma, L. & Niu, J. THREE□DIMENSIONAL (3D) ANTHROPOMETRY AND ITS APPLICATIONS IN PRODUCT DESIGN. Handbook of human factors and ergonomics, 281–302 (2021).

2 Niu, J. & Li, Z. Using three-dimensional (3d) anthropometric data in design. Handbook of anthropometry: physical measures of human form in health and disease, 3001–3013 (2012).

3 Bruneau, D. A. & Cronin, D. S. Head and neck response of an active human body model and finite element anthropometric test device during a linear impactor helmet test. Journal of Biomechanical Engineering 142, 021004 (2020).

4 Mandolini, M., Caragiuli, M., Brunzini, A., Mazzoli, A. & Pagnoni, M. A procedure for designing custom-made implants for forehead augmentation in people suffering from Apert syndrome. Journal of Medical Systems 44, 1–10 (2020).

5 Sena, K. & Piyasin, S. Determination of average contour of Thais skulls for design of implants. American J of Engineering and Applied Sciences 1, 168–173 (2008).

6 Dong, Y., Zhao, Y., Bai, S., Wu, G. & Wang, B. Three-dimensional anthropometric analysis of the Chinese nose. Journal of plastic, reconstructive & aesthetic surgery 63, 1832–1839 (2010).

7 Rodriguez-Florez, N. et al. Statistical shape modeling to aid surgical planning: associations between surgical parameters and head shapes following spring-assisted cranioplasty. International journal of computer assisted radiology and surgery 12, 1739–1749 (2017).

8 Wang, J., Thornton, J., Kolesnik, S. & Pierson Jr, R. Anthropometry in body composition: an overview. Annals of the New York Academy of Sciences 904, 317–326 (2000).

9 Boehnen, C. & Flynn, P. in Fifth International Conference on 3-D Digital Imaging and Modeling (3DIM’05). 310-317 (IEEE).

10 Bougourd, J., Treleaven, P. & Allen, R. in Eurasia-Tex Conference, Donghua University, Shanghai, China.

11 Isaacs, M. 3D fit for the future-3D fit and design can be the solution to several industry problems. AATCC Review-American Association of Textile Chemists and Colorists 5, 21–24 (2005).

12 Seidl, A., Trieb, R. & Wirsching, H.-J. in Proceedings of 17th World Congress on Ergonomics.

13 Luximon, Y., Martin, N. J., Ball, R. & Zhang, M. Merging the point clouds of the head and ear by using the iterative closest point method. International Journal of the Digital Human 1, 305–317 (2016).

14 Zhuang, Z., Shu, C., Xi, P., Bergman, M. & Joseph, M. Head-and-face shape variations of US civilian workers. Applied ergonomics 44, 775–784 (2013).

15 Skals, S., Ellena, T., Subic, A., Mustafa, H. & Pang, T. Y. Improving fit of bicycle helmet liners using 3D anthropometric data. International Journal of Industrial Ergonomics 55, 86–95 (2016).

16 Luximon, Y., Ball, R. & Justice, L. The 3D Chinese head and face modeling. Computer-Aided Design 44, 40–47 (2012).

17 Perret-Ellena, T., Skals, S. L., Subic, A., Mustafa, H. & Pang, T. Y. 3D anthropometric investigation of head and face characteristics of Australian cyclists. Procedia Engineering 112, 98–103 (2015).

18 Luximon, A., Zhang, Y., Luximon, Y. & Xiao, M. Sizing and grading for wearable products. Computer-Aided Design 44, 77–84 (2012).

19 Lacko, D. et al. Product sizing with 3D anthropometry and k-medoids clustering. Computer-Aided Design 91, 60–74 (2017).

20 Verwulgen, S. et al. A new data structure and workflow for using 3D anthropometry in the design of wearable products. International Journal of Industrial Ergonomics 64, 108–117 (2018).

21 Huang, Y. et al. A Morphometric Study of Head in the Population of Young Chinese Adults. Int. J. Morphol 1, 2 (2023).

22 Zhang, J., Iftikhar, H., Shah, P. & Luximon, Y. Age and sex factors integrated 3D statistical models of adults’ heads. International Journal of Industrial Ergonomics 90, 103321 (2022).

23 Ellena, T., Skals, S., Subic, A., Mustafa, H. & Pang, T. Y. 3D digital headform models of Australian cyclists. Applied ergonomics 59, 11–18 (2017).

24 Ellena, T., Subic, A., Mustafa, H. & Yen Pang, T. A novel hierarchical clustering algorithm for the analysis of 3D anthropometric data of the human head. Computer-Aided Design and Applications 15, 25–33 (2018).

25 Kuo, C.-C., Wang, M.-J. & Lu, J.-M. Developing sizing systems using 3D scanning head anthropometric data. Measurement 152, 107264 (2020).

26 Niu, J., Li, Z. & Salvendy, G. Multi-resolution shape description and clustering of three-dimensional head data. Ergonomics 52, 251–269 (2009).

27 Seo, H., Kim, J. I. & Kim, H. Development of Korean head forms for respirator performance testing. Safety and Health at Work 11, 71–79 (2020).

28 Zhang, J., Zhou, K., Luximon, Y., Li, P. & Iftikhar, H. 3D-guided facial shape clustering and analysis. Multimedia Tools and Applications 81, 8785–8806 (2022).

29 Ray, S. & Turi, R. H. in Proceedings of the 4th international conference on advances in pattern recognition and digital techniques. 143 (Calcutta, India).

30 Rousseeuw, P. J. Silhouettes: a graphical aid to the interpretation and validation of cluster analysis. Journal of computational and applied mathematics 20, 53–65 (1987).

31 Nainggolan, R., Perangin-angin, R., Simarmata, E. & Tarigan, A. F. in Journal of Physics: Conference Series. 012015 (IOP Publishing).

32 Osborne, J. W. & Overbay, A. The power of outliers (and why researchers should always check for them). *Practical Assessment*, Research, and Evaluation 9, 6 (2019).

33 Lacko, D. et al. Evaluation of an anthropometric shape model of the human scalp. Applied ergonomics 48, 70–85 (2015).

34 Heutinck, P. et al. Statistical shape modeling for the analysis of head shape variations. Journal of Cranio-Maxillofacial Surgery 49, 449–455 (2021).

35 Zhang, J., Luximon, Y., Shah, P., Zhou, K. & Li, P. Customize my helmet: A novel algorithmic approach based on 3D head prediction. Computer-Aided Design 150, 103271 (2022).

36 Piegl, L. & Tiller, W. The NURBS book. (Springer Science & Business Media, 1996).

37 Amor, B. B., Ardabilian, M. & Chen, L. in *Third International Symposium on 3D Data Processing*, Visualization, and Transmission (3DPVT’06). 279–286 (IEEE).

38 Xi, W., Zongqian, W. & Qiao, L. Extraction of Feature Points for Non-Uniform Rational B-Splines (NURBS)-Based Modeling of Human Legs. Journal of Donghua University 39 (2022).

39 Xie, S., Li, N. & Lv, Z. Human Head Modeling Using NURBS Method. Advances in Neural Network Research and Applications, 479–484 (2010).

40 Pal, P. & Ballav, R. Object shape reconstruction through NURBS surface interpolation. International journal of production research 45, 287–307 (2007).

41 Zhang, B. & Molenbroek, J. Representation of a human head with bi-cubic B-splines technique based on the laser scanning technique in 3D surface anthropometry. Applied ergonomics 35, 459–465 (2004).

42 Hadi, N. A. & Alias, N. in Proceedings of the 2019 4th International Conference on Intelligent Information Technology. 16–20.

43 Hsiao, H. et al. Comparison of measured and self-reported anthropometric information among firefighters: implications and applications. Ergonomics 57, 1886–1897 (2014).

44 Hsiao, H. et al. Sizing firefighters: method and implications. Human factors 56, 873–910 (2014).

45 Hsiao, H., Whisler, R. & Bradtmiller, B. Needs and procedures for a national anthropometry study of law enforcement officers. Human Factors 65, 403–418 (2023).

46 Robinette, K. M., et al. Civilian American and European surface anthropometry resource (CAESAR), final report, volume I: Summary. Sytronics Inc Dayton Oh (2002).

47 Zhuang, Z. & Bradtmiller, B. Head-and-face anthropometric survey of US respirator users. Journal of occupational and environmental hygiene 2, 567–576 (2005).

48 Ball, R. M. SizeChina: A 3D anthropometric survey of the Chinese head. (2011).

49 Wang, M.-J. J., Wang E. M.-y. & Lin, Y.-C. The anthropometric database for children and young adults in Taiwan. Applied Ergonomics 33, 583–585 (2002).

50 Molenbroek, J., Albin, T. J. & Vink, P. Thirty years of anthropometric changes relevant to the width and depth of transportation seating spaces, present and future. Applied ergonomics 65, 130–138 (2017).

51 Malkoc, İ., et al. Are New Generations Getting Bigger in Size? Anthropometric Measurements in Erzurum. The Eurasian Journal of Medicine 46, 192 (2014).

52 NIOSH. NIOSH Anthropometric Data and ISO Digital Headforms, <https://www.cdc.gov/niosh/data/datasets/rd-10130-2020-0/default.html> (April 16, 2020).

53 Gordon, C. C. et al. 2012 anthropometric survey of us army personnel: Methods and summary statistics. Army Natick Soldier Research Development and Engineering Center MA, Tech. Rep (2014).

54 Akoglu, H. User’s guide to correlation coefficients. Turkish journal of emergency medicine 18, 91–93 (2018).

55 MacQueen, J. in Proceedings of the fifth Berkeley symposium on mathematical statistics and probability. 281-297 (Oakland, CA, USA).

56 Kaufman, L. & Rousseeuw, P. J. Finding groups in data: an introduction to cluster analysis. (John Wiley & Sons, 2009).

57 Weisberg, S. Applied linear regression. Vol. 528 (John Wiley & Sons, 2005).

58 Ostertagová, E. Modeling using polynomial regression. Procedia Engineering 48, 500–506 (2012).

59 Harmening, C. & Neuner, H. in Proceedings of the 3rd Joint international Symposium on Deformation Monitoring (JISDM), Vienna, Austria.

60 Paquette, S. *Anthropometric survey (ANSUR) II pilot study: methods and summary statistics*. (Anthrotch, US Army Natick Soldier Research, Development and Engineering Center, 2009).

61 Zhuang, Z., Landsittel, D., Benson, S., Roberge, R. & Shaffer, R. Facial anthropometric differences among gender, ethnicity, and age groups. Annals of occupational hygiene 54, 391–402 (2010).

62 Gordon, C. C. 1988 Anthropometric survey of US army personnel: methods and summary statistics. Technical Report Natick/TR-89/044 (1989).

63 Zhuang, Z., Benson, S. & Viscusi, D. Digital 3-D headforms with facial features representative of the current US workforce. Ergonomics 53, 661–671 (2010).

64 McFadden, D. Conditional logit analysis of qualitative choice behavior. (1973).

65 Goleij, N., Hafezi, P. & Ahmadi, O. Investigating the trends and causes of changes in human anthropometric dimensions over the past three decades: a challenge for ergonomic design. International Journal of Occupational Safety and Ergonomics 30, 480–485 (2024).

66 Taylor, N. A., Fullagar, H. H., Mott, B. J., Sampson, J. A. & Groeller, H. Employment standards for Australian urban firefighters: Part 1: The essential, physically demanding tasks. Journal of occupational and environmental medicine 57, 1063–1071 (2015).

67 Henderson, N. D. Predicting long□term firefighter performance from cognitive and physical ability measures. Personnel Psychology 63, 999–1039 (2010).

68 Hsiao, H., Long, D. & Snyder, K. Anthropometric differences among occupational groups. Ergonomics 45, 136–152 (2002).

69 Henneberg, M. & Ulijaszek, S. J. Body frame dimensions are related to obesity and fatness: lean trunk size, skinfolds, and body mass index. American Journal of Human Biology: The Official Journal of the Human Biology Association 22, 83–91 (2010).

70 Kwak, S. G. & Kim, J. H. Central limit theorem: the cornerstone of modern statistics. Korean journal of anesthesiology 70, 144 (2017).

71 Madhulatha, T. S. in International Conference on Advances in Computing and Information Technology. 472-481 (Springer).

72 Velmurugan, T. & Santhanam, T. Computational complexity between K-means and K-medoids clustering algorithms for normal and uniform distributions of data points. Journal of computer science 6, 363 (2010).

73 Batra, A. in ICACCT, 5th IEEE International Conference on Advanced Computing & Communication Technologies. 274-279.

74 Kuželka, K. & Marušák, R. Comparison of selected splines for stem form modeling: A case study in Norway spruce. Annals of Forest Research, 137–148 (2014).

75 Lahtinen, A. On the construction of monotony preserving taper curves. (1988).

76 Ma, W. & Kruth, J.-P. NURBS curve and surface fitting for reverse engineering. The International Journal of Advanced Manufacturing Technology 14, 918–927 (1998).

77 Carlson, N. (2009).

78 Piegl, L. & Tiller, W. Curve and surface constructions using rational B-splines. Computer-aided design 19, 485–498 (1987).

79 Dimas, E. & Briassoulis, D. 3D geometric modeling based on NURBS: a review. Advances in Engineering Software 30, 741–751 (1999).

80 Piegl, L. On NURBS: a survey. IEEE Computer Graphics and Applications 11, 55–71 (1991).

81 Sinthanayothin, C. & Bholsithi, W. in 2009 6th International Conference on Electrical Engineering/Electronics, Computer, Telecommunications and Information Technology. 668–671 (IEEE).

82 Penguian. NURBS Toolbox by D.M. Spink (https://www.mathworks.com/matlabcentral/fileexchange/26390-nurbs-toolbox-by-d-m-spink), MATLAB Central File Exchange. (2024).

